# EV Fingerprinting: Resolving extracellular vesicle heterogeneity using multi-parametric flow cytometry

**DOI:** 10.1101/2022.11.10.515864

**Authors:** Ariana K. von Lersner, Fabiane C. L. Fernandes, Patricia M. M. Ozawa, Sierra M. Lima, Tatyana Vagner, Bong Hwan Sung, Mohamed Wehbe, Kai Franze, John T. Wilson, Jonathan M. Irish, Alissa Weaver, Dolores Di Vizio, Andries Zijlstra

**Affiliations:** Program in Cancer Biology, Vanderbilt University, Nashville, TN, USA; The Center for EV Research, Vanderbilt University, Nashville, TN, USA; Department of Pathology, Microbiology and Immunology, Vanderbilt University Medical Center, Nashville, TN, USA; Institute of Applied Biosciences and Chemistry, Hogeschool Arnhem en Nijmegen University of Applied Sciences, Nijmegen, Netherlands; Department of Cell and Developmental Biology, Vanderbilt University School of Medicine, Nashville, TN, USA; Department of Surgery, Cedars-Sinai Medical Center, Los Angeles, USA; Department of Chemical and Biomolecular Engineering, Vanderbilt University, Nashville, TN, USA; KNIME GmbH, Konstanz, Germany; Department of Research Pathology, Genentech Research and Early Development, San Francisco, CA, USA

## Abstract

Mammalian cells release a heterogeneous array of extracellular vesicles (EVs) that impact human biology by contributing to intercellular communication. To resolve EV heterogeneity and define the EV populations associated with specific biological processes, we developed a method named “EV Fingerprinting” that discerns distinct vesicle populations using dimensional reduction of multi-parametric data collected by quantitative single-EV flow cytometry. After validating this method against synthetic standards, the EV Fingerprinting analysis of highly purified EVs enabled a much more granular resolution of biochemically distinct EV populations than previously established methods. The analysis of EVs produced after molecular perturbation of EV biogenesis through ablation of the GTPase Rab27a and overexpression of the tetraspanin CD63 revealed that EV Fingerprinting reflects the molecular state of a cell. Subsequent analysis of human plasma demonstrates the capacity of EV Fingerprinting to resolve EV populations in complex biological samples and detect tumor-cell derived EVs.

## Introduction

Extracellular vesicles (EVs) are functional contributors to intercellular communication in health and disease^1^. The detection and characterization of these vesicles is therefore thought to be a means to derive biological insights and attain biomarkers that inform on the status of a patient. Although frequently referred to as a single class, cumulatively defined as lipid bilayer enclosed particles, the term “extracellular vesicles” encompasses a heterogeneous group of particles that originate through multiple biogenesis pathways and vary greatly in both size and composition^2^. EVs are commonly characterized by their size. While imperfect, this metric is informative in distinguishing between smaller EVs (S-EV) ranging from 30 nm – 200 nm and larger EVs (L-EV) ranging from 200 nm to ≥ 1000 nm.

While much remains unknown about EV biogenesis, it has been established that they are generated through at least two distinct mechanisms (Figure 1a): i) Cells can produce EV through exocytosis of endosome-derived multivesicular bodies (MVBs) that fuse with the plasma membrane to release exosomes into the extracellular space. This distinct biogenesis pathway is regulated by the GTPase Rab27a which controls docking of MVBs with the plasma membrane. ^34^ These EVs are thought to be ≤ 200 nm. ii) Alternatively, EVs can arise through direct budding and fission of the plasma membrane (ectocytosis) to produce ectosomes. These vesicles range broadly in size and can be as small as the exosomes but also significantly larger (≥1µm).^5,6^

**Figure 1:**
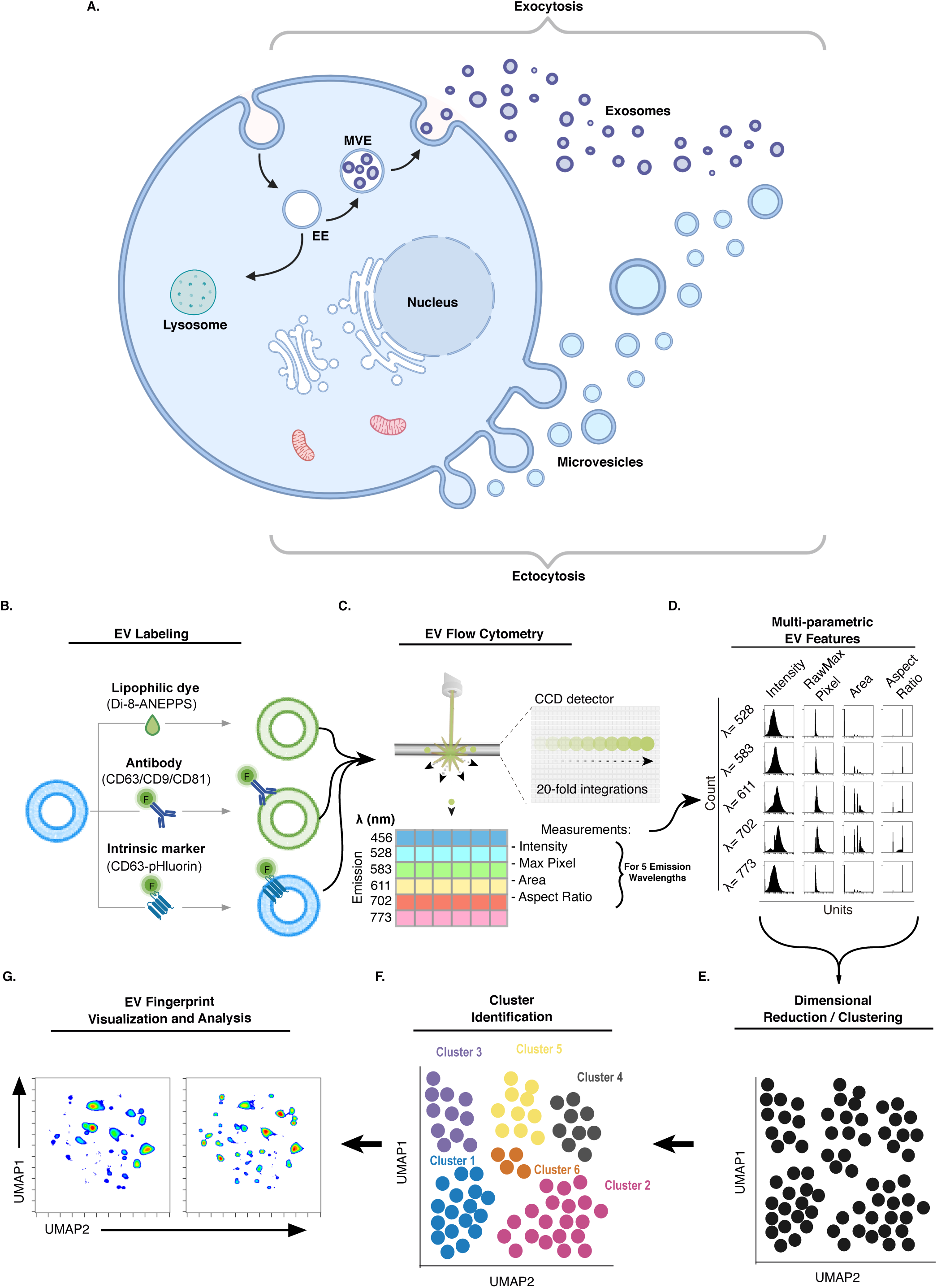
EV Fingerprinting overview. **a,** A representation of EV biogenesis through exocytosis and ectocytosis. Created with BioRender.com. **b,** Extrinsic labeling of EVs using the lipophilic dye di-8-ANEPPS (di8) and fluorescently conjugated antibody or intrinsic labeling using fluorescent fusion proteins (pHluorin_M153R tagged CD63). **c,** Optical triggering on the CellStream™ uses a time delay integration charge-coupled device (TDI-CCD) to collect measurements (including Intensity, RawMax Pixel, Area, Aspect Ratio) simultaneously for up to six emission wavelengths (em = 456, 528, 583, 611, 702, and 773). **d,** A matrix histogram of measurements for the five emission wavelengths collected for di8 (excitation = 488) demonstrates the multi-parametric nature of EV features. **e,** Dimensional reduction embedding and clustering of multi-parametric EV features using UMAP. **f,** Cluster identification using HDBSCAN. **g,** Representative results of a UMAP density plot comparing two paired samples.

Not only do EVs vary in size, they are also known to vary in lipid, protein, carbohydrate, and nucleic acid composition^7–10^. These variations in geometry, chemical composition, and molecular cargo underlie the heterogeneity of EVs produced by living systems^11^. Given that EVs have been demonstrated to convey a diverse array of biological activity,^12^ the heterogeneity of EV populations suggests that unique populations of EVs convey distinct functions. This is underscored by studies evaluating the functional role of S-EV vs. L-EV^13–15^. Moreover, EV heterogeneity can reflect a biological state (i.e. cancer vs normal) thereby serving as biomarkers.^16,17^ Given the complexity of EV biogenesis, we hypothesize that resolving EV heterogeneity using dimensional reduction of multi-parametric measurements will yield an “EV Fingerprint” unique to the biological state of the cell, tissue, or organism from which it was derived.

Single-EV flow cytometry methods are gaining popularity given their ability to characterize molecular cargoes on a single vesicle basis. While most conventional flow cytometers are not set up to detect particles smaller than 500 nm in diameter, instrumentation can be adapted to enable small particle detection.^18^ For example, Stoner et. al.^19^ built a customized high-sensitive flow cytometer to extend the limit of detection to 70-80 nm. In addition, fluorescence triggering with lipophilic dyes such as with di-8-ANEPPS (di8) demonstrated reproducible detection of EVs^19^. Other specialized single-EV flow cytometry platforms achieve increased instrument sensitivity, in part by slowing down the flow rate which, in turn, greatly limits throughput (10,000 - 12,000 events/min)^18,20^. Conversely, technologies with faster flow rates (≥100,000 events/min) not only suffer from lower sensitivity, but are also susceptible to the so-called swarm effect, in which groups of small particles are registered as single large particles^21,22^.

To achieve accurate high-throughput detection of small particles at a fast flow rate (~100,000 events/min), the Amnis CellStream™ platform utilizes a Time Delay Integration Charge-Coupled Device (TDI-CCD). The detection of photons across spatially separated pixels permits the capture of multi-parametric image-like features, providing new data dimensions. We exploited the gain in detection sensitivity with the pixel-based image features to resolve EV heterogeneity.

In this study we leverage an optimized staining strategy using the lipophilic dye di8, coupled with dimensional reduction of the new data dimensions collected by a TDI-CCD, to develop a novel “EV Fingerprinting” approach that deconvolves the complexity of EV populations in biological samples. The method was established with highly purified EV standards and validated with orthogonal EV characterization methods by leveraging the guidelines from the International Society for Extracellular Vesicles (ISEV) for the minimal information for studies of EVs (MISEV)^23–25^. EV Fingerprinting was subsequently implemented to reveal the existence of unique EV populations that: i) can be parsed by size using differential centrifugation, ii) differentially impacted by disruption of exosome secretion with knockdown (KD) of Rab27a, and iii) uniquely impacted by the overexpression of EV cargo (CD63). Finally, the method was successfully implemented to quantitatively detect, resolve, and characterize EV populations in plasma.

## Results

### Principle of EV Fingerprinting and Method Overview

In order to develop a method to deconvolve the heterogeneity of EVs, we used single-EV detection by flow cytometry after fluorescent labeling followed by dimensional reduction and machine learning on multiparametric optical features (Figure 1b-g). The approach leverages changes in fluorescence intensity and emission spectra of the lipophilic dye di8, associated with variations in particle size and lipid composition, to detect, discern, and quantify distinct EV populations. In addition, single-EV cargoes are visualized using a genetically encoded fluorescent marker (e.g. pHluorin-CD63) ^26^ or a cargo-specific antibody (e.g. anti-CD63 antibody, Fig. 1b). Fluorescently labeled EVs are detected in the Amnis Cellstream™ flow cytometer, where 20-fold signal integration of six potential fluorescent emissions collected by the TDI-CCD detector enables sensitive detection of passing particles (Fig. 1c). Multi-parametric data from di8 staining is extracted from the final integration using four optical features across five emission wavelengths (Fig. 1d, Supplementary Table 1 and 2). Data reduction using Uniform Manifold Approximation Projection (UMAP) of the resulting 20 dimensions creates a 2D representation of the EV populations present in the sample (Fig. 1e). Hierarchical Density-Based Spatial Clustering of Applications with Noise (HDBSCAN) is subsequently used for cluster identification (Fig. 1f)^27–30^. The resulting pattern of clusters is a “fingerprint” of distinct EV populations. Unlike previously established bulk EV analytical methods, a quantitative assessment of clusters within the EV Fingerprint makes it possible to determine how experimental manipulation, molecular perturbation, or disease state alters select EV populations (Fig. 1g).

### Optimization of EV detection by fluorescent triggering

Single-EV detection was achieved using optimized di8 staining of EVs followed by fluorescence-triggered flow cytometry (Amnis Cellstream™) using a TDI-CCD detector to collect optical measures upon fluorescence excitation by a 488 laser. To enable the analysis of both unpurified and purified EV samples across large variations in EV concentrations, two staining methods were established to enable analysis of samples with either low or high particle concentrations (Methods #1 and #2 in Extended Data. Fig 1a and 1b, respectively). Method #1: For unpurified, EV-containing fluids, the sample is stained with a low concentration of di8 (0.25 µM) and analyzed directly by flow cytometry. Method #2: For purified EVs, the sample is stained with a high concentration of di8 (2 µM). The stained EVs are subsequently diluted 200-fold before analysis by flow cytometry. This dilution reduces background fluorescence and avoids swarming the detector with a high number of particles.^31,32^ Serial dilution is frequently implemented to ensure detection within the quantitative range of the assay. These methods leverage the increase in quantum efficiency when di8 transitions from the aqueous buffer to the hydrophobic lipid bilayer^33^. To identify the optimal staining conditions, a series of trials was completed to evaluate the impact of: 1) di8 concentration (Extended Data Fig. 1a-d), 2) the staining temperature (Extended Data Fig. 1d), 3) and the laser excitation intensity (Extended Data Fig. 1e, f). Based on the following observations, we defined the optimal staining conditions as summarized in Supplementary Table 3.

Using Method #1, we tested EV detection across a serial dilution of di8 from 4 µM to 0.0078 µM staining at room temperature (RT). Consistent EV detection was observed in a range from 0.0156-0.25µM (Extended Data Fig. 1c, 100K EV-blue) with minimal signal derived from free dye (Extended Data Fig. 1c, Buffer-black).

While lipophilic labeling is typically performed at RT, antibody staining for flow cytometry is often optimized at different temperatures. To assess the impact of temperature on vesicle staining, Method #2 was evaluated at RT and 37°C using 2-fold dilutions from 4 µM to 0.5 µM. EV detection was consistent for both temperatures at 2 µM di8 (Extended Data Fig. 1b and d).

The impact of excitation energy on EV detection was assessed using a 488 nm laser titration for both Method #1 and #2. EV detection plateaued at laser powers greater than 45% for both staining methods (Extended Data Fig. 1e,f). Moreover, background particle detection in the absence of EVs increased when laser power was increased beyond 25% with both staining methods (Extended Data Fig. 1e,f).

### Quantitative single-EV detection

Single-EV detection in a flow cytometer can require extensive dilution of the source material to avoid swarming the detector with multiple particles. However, the resulting reduction in molecular density is known to impact both microparticles and macromolecules by promoting protein:lipid and protein:protein interactions^34^. To offset the diminution in protein and other buffering components caused by sample dilution,^34^ we tested the benefit of molecular crowding (MC) with dextran (Mr = ~100,000). Final dextran concentrations of 1.625%, 3.25%, and 6.5% compared to PBS (0% Dextran) improved single particle detection for both FITC-labeled nano-sized beads as well as UC purified di8-stained EVs by 100-400% while increasing dextran concentration to 10% eliminated gains in detection (Extended Data Fig. 2a,b respectively). MC at 3.25% dextran was subsequently used as the optimal condition for particle detection (Extended Data Fig. 2c,d).

Single particle detection was verified using non-fluorescent and fluorescent synthetic bead standards as well as UC purified di8-stained EVs (Extended Data Fig. 2c,d). FITC-labeled sizing beads (0.5, 0.2, and 0.1 µm respectively) were serially diluted and the corresponding linear detection was successfully quantified (Extended Data Fig. 2c). To define the quantitative range of EV detection, 100K UC purified EVs stained by Method #2 were evaluated across an extended serial dilution range (Extended Data Fig. 2d). The linear range of EV detection was identified at 50,000-400,000 EV/µl (~2.5-22*10^6 EV/mL) for which the EV count corresponded linearly to the dilution factor and the median fluorescence intensity (MFI, Median 488-611) was stable (grey box, Extended Data Fig. 2d). The specificity of lipid-based vesicle detection was confirmed through detergent lysis of the EVs. Indeed, detection of di8-positive (di8+) EVs was abrogated upon lysis with 0.5% NP40 detergent, confirming specificity of EV detection (Extended Data. Fig. 2e).

The validated EV flow cytometry strategy was subsequently applied to EV preparations generated by a standard sequential UC method^35^ designed to enrich for L-EVs at 10,000 x g (10K) and for S-EVs at 100,000 x g (100K) (Fig. 2). TEM confirmed the expected enrichment in the relative fractions (Fig. 2a, scale bar = 200 nm). In parallel, nanoparticle tracking analysis (NTA) confirmed the differential enrichment for larger EVs in the 10K UC prep and for smaller EVs in the 100K UC prep (Fig. 2b). Flow cytometry successfully detected both intrinsically labeled EVs (pHluorin-CD63) and extrinsically labeled EVs (di8) from UC purified EV preparations (10K and 100K) as well as from unfractionated conditioned medium (CM) (Fig. 2c). The lower MFI and scatter measurements from 100K vs. the 10K preparations is consistent with the reduced size of the EVs in 100K preps observed by TEM and NTA.

**Figure 2:**
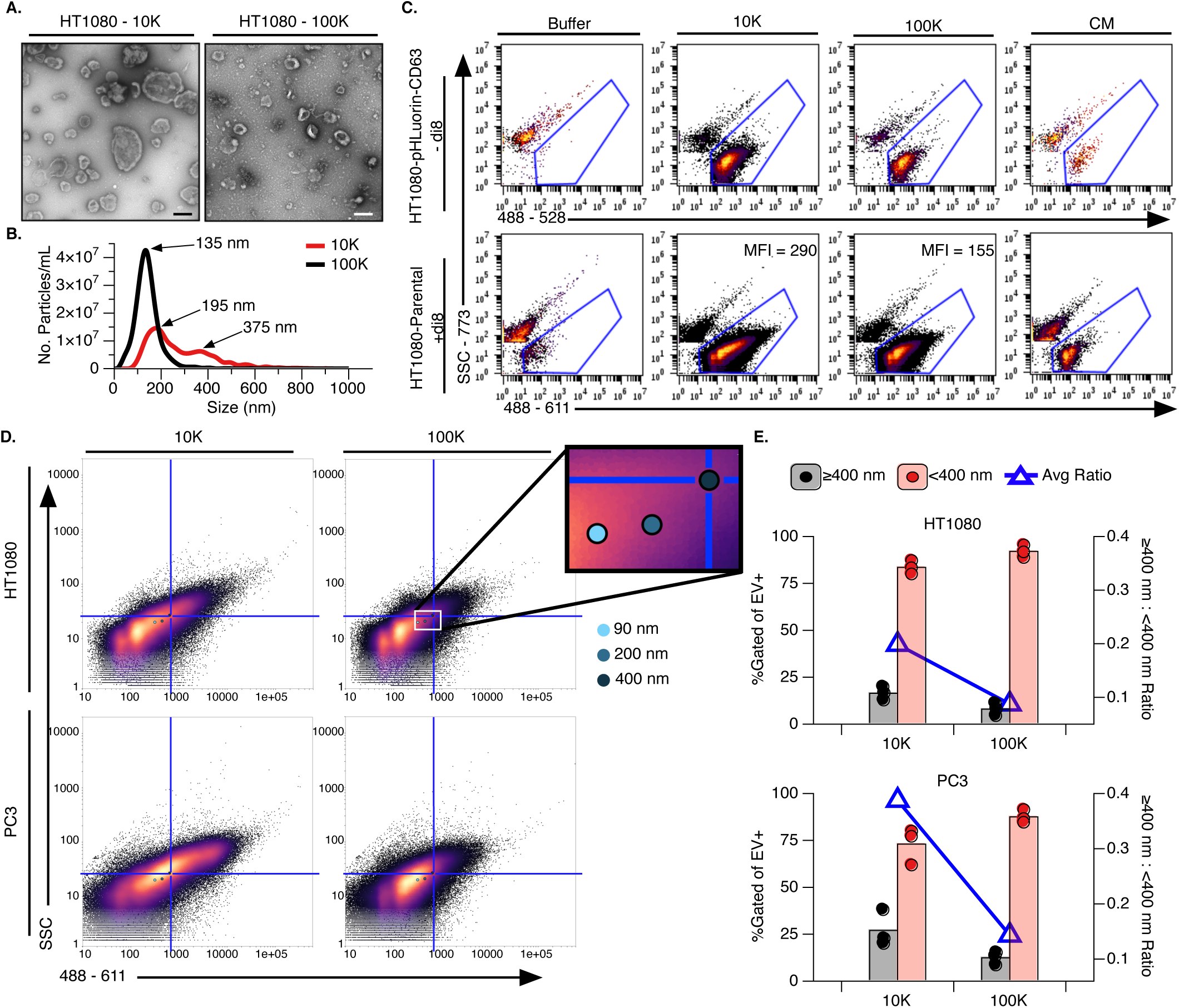
Quantitative detection of single EVs by flow cytometry. **a,** Negative staining of transmission electron micrographs (TEM) of 10K (left) and 100K (right) UC purified EV preps from HT1080 parental cells. Scale bar, 200 nm. **b,** Representative traces from nanoparticle tracking analysis (NTA) of 10K and 100K EV preparations. **c,** Representative flow cytometry scatter plots of intrinsically labeled HT1080-pHluorin-CD63 (top) and extrinsically di-8-ANEPPS (di8) stained HT1080-Parental (bottom) samples from 10K and 100K fresh d-UC purified EV preps and conditioned media (CM). EV+ gating was performed against the buffer controls. **D,** EV+ events from 10K and 100K UC purified EVs from HT1080-Parental and PC3 cells. The median SSC and 488-611 (excitation - emission) MFI of sizing liposomes (90 nm, 200 nm, and 400 nm) is shown within the scatter plots, as indicated (from Extended Data Fig. 3). Sizing liposome data were measured in triplicate. The upper right quadrant represents EVs ≥400 nm. **e,** Production of EVs ≤ (pink) or ≥ (grey) 400 nm from HT1080 and PC3 cell lines quantified across three serial dilutions. Bar plots represent quantitation of the dilution-corrected average EV+ events ≥400 nm (grey, upper right quadrant on **(d)**) and ≤400 nm (pink, lower left quadrant on **(d)**) on the left y-axis. Individual dilution-corrected replicates are shown as points. The average ratio of larger to smaller EV is shown on the right y-axis (blue triangle).

To assess the ability of flow cytometry to reveal size-based enrichment of EVs isolated by differential UC (d-UC), EVs were mapped to sizing liposomes (90 – 400 nm) labeled with the same di8-staining Method #2 (Extended Data Fig. 3). Given that PC3 cells have been previously characterized to shed L-EVs including Large Oncosomes (LO, ≥1 µm)^36,37^, we used the largest sizing liposome (400 nm) as a landmark to evaluate enrichment of S-EV (<400 nm) and L-EV (≥400 nm) populations across the UC preparations (10K vs 100K, Fig. 2d,e). Plotting the MFI of sizing liposome standards over scatter plots of 10K and 100K preparations from two cell lines (fibrosarcoma (HT1080) and prostate cancer (PC3)) illustrates that our single EV flow cytometry method could indeed detect EVs across a large size range (Fig. 2d,e). Consistent with the NTA observations (Fig. 2b), this method detected a higher frequency of smaller EVs (<400 nm) in both 10K and 100K UC preparations, but a distinct enrichment (blue line) of larger EVs (≥400 nm) was observed in 10K UC preparations (Fig. 2e). This enrichment is 2-fold higher for PC3 than HT1080 cells, consistent with previously published reports (Fig. 2e).^36^

### EV Fingerprinting

During single-EV detection on the Amnis CellStream™, the TDI-CCD camera collects 14 optical features for each of the 6 emission channels (total = 84 features) for every passing particle (Fig. 1c-d, Supplementary Tables 1 and 2). These optical EV features were leveraged in a multiparametric analysis to discriminate between distinct EV populations (See Fig. 1 for summary).

In principle, di8 intensity should increase with lipid mass (EV size). In addition, di8 has a broad emission spectrum which shifts with changes in lipid composition of the lipid bilayer it occupies^38–40^. Thus, the spectral emission profile of di8-labeled EVs changes when the lipid composition (e.g. cholesterol) is altered (Extended Data Fig. 3a). Indeed, using 90-400 nm sizing liposomes made of the same lipid composition (Supplementary Table 4), we observed that the fluorescence intensity of di8 stained liposomes increases as a function of liposome size (excitation 488 nm, emission 611 nm, Extended Data Fig. 3b,c). Conversely, changing the cholesterol content of the liposomes while keeping their size constant (100 nm, Supplementary Table 5) revealed a distinct spectral emission shift from 702 nm for liposomes with low cholesterol content to 611 nm for liposomes with high cholesterol content without significantly altering the total fluorescence intensity (Extended Data Fig. 3d and 3e, respectively). This relationship in emission shift is subsequently quantified using the 611:702 emission ratio (Extended Data Fig. 3f). Thus, MFI can be used as a proxy for size while the 611:702 emission ratio can be leveraged to inform on lipid composition (Extended Data Fig. 3).

We subsequently used dimensional reduction and cluster analysis to deconvolve the heterogenous EVs detected by single EV flow cytometry into distinct populations. This allowed us to determine which EV populations were differentially enriched during UC purification, and which EV populations were impacted by molecular perturbation of EV biogenesis pathways (Fig. 3, 4, and 5). The analysis workflow is comprised of ingesting raw .fcs files, followed by dimensional reduction of arcsinh transformed features, and subsequent visualization in the online flow cytometry software, Cytobank (Extended Data Fig. 4a). To facilitate the analysis, a two-phase data processing, analysis, and visualization workflow was developed in the data science platform KNIME (Extended Data Fig. 4b, see methods)^41,42^. During the first phase (upper panel), a portion of flow cytometry events from each sample is arcsinh transformed, reduced in dimension using UMAP, and clustered using HDBSCAN based on 20 di8 features (Supplementary Tables 2 and 6). In the second phase (bottom panel), select HDBSCAN clusters are filtered based on user input parameters and re-examined with the same features or with additional features such as those derived from antibody fluorescent channels (Supplementary Tables 2 and 6). In this study, re-examination was used for two purposes: 1) data cleaning: to remove non-specific particles, such as those found in the control buffers or cell-free CM in preparation for downstream evaluations (Extended Data Fig. 4b) and 2) focused cluster analysis: to re-examine an individual cluster with a new set of features, such as the detection of a cargo (e.g. anti-CD63).

To determine if EV Fingerprinting could be used to quantitatively assess heterogeneous populations, the method was used to analyze fluorescent sizing beads of known quantity and to compare with manual gating (Extended Data Fig. 5). Indeed the analysis generated bead-specific clusters and distinguished them from background particles (Extended Data Fig. 5a,b). Quantitation was accurate for both individual bead sizes as well as for a mixture of all three sizes Extended Data Fig. 5c).

EV Fingerprinting was subsequently applied to EVs labeled intrinsically with fluorescently-tagged CD63 EV-reporter (pHluorin-CD63)^26^ or extrinsically with di8. Flow cytometry of unpurified CM and EVs isolated by sequential 10K and 100K UC was used to evaluated the ability of EV Fingerprinting to assess the enrichment of EV from culture medium by ultracentrifugation. The quantitative range for EV flow cytometry was determined by serial dilution, as shown in Extended Data Figure 2d. Intrinsically (pHluorin-CD63) and extrinsically (di8) labeled EV samples were analyzed using the dimensional reduction workflow (Extended Data Fig. 6). It was possible to observe EV populations differentially enriched between 10K and 100K UC preparations in both intrinsically and extrinsically labeled EVs (10K vs 100K, Extended Data Fig. 6). However, the dimensional reduction of di8-labeled EVs revealed a much more detailed array of populations than pHluorin-CD63 EVs (Extended Data Fig. 6, -di8 HT1080-pHluorin-CD63 vs +di8 HT1080-Par). Consequently, subsequent studies leveraged only extrinsic di8 labeling.

### EV Fingerprinting captures size-based enrichment of EV populations by density gradient UC purified EV

EVs are commonly isolated by sequential UC at increasing speeds (2.8K, 10K, and 100K) followed by centrifugal upward flotation in a density gradient (DG-UC)^35,43^. This method leverages the size and density of EVs to separate them from other lipid-containing particles and proteins. Moreover, DG-UC is a common method to enrich for S-EVs at 100K, while L-EVs are enriched at 2.8-10K^37^.

We tested the capacity of EV Fingerprinting to detect the differential enrichment of EV populations by DG-UC at 2.8K (2K), 10K, and 100K from PC3 cells (Fig. 3). Tunable resistive pulse sensing (TRPS) sizing measurements of the isolated EV preparations from PC3 cells, known to secrete both L-EVs and S-EVs,^37^ confirmed larger vesicles (> 200 nm) isolated in the 2K and 10K preparations compared to the 100K preparation (Extended Data Fig. 7a-c). Flow cytometry of di8-labeled EV preparations readily detected EVs in all three preparations when compared to buffer alone (Extended Data Fig. 7d). The detection of a low number of particles with high MFI in the 2K preparation compared to high number of particles with lower MFI in 100K is consistent with the TRPS observation that the slower speed preparations contain fewer but larger (and thus brighter) particles, while high speed preparations contain more but smaller (and thus dimmer) particles. By comparison, the 10K preparation seems to contain a mixture of both larger and smaller EVs. EV Fingerprinting of DG-UC purified EVs reveals over 80 distinct populations (Extended data 7e). After removal of buffer-derived clusters and rank-ordering the populations according to abundance (Extended Data Fig. 7f), clusters containing ≥ 1% of total data were re-examined for differential enrichment between 2K, 10K and 100K preparations (Fig. 3).

**Figure 3:**
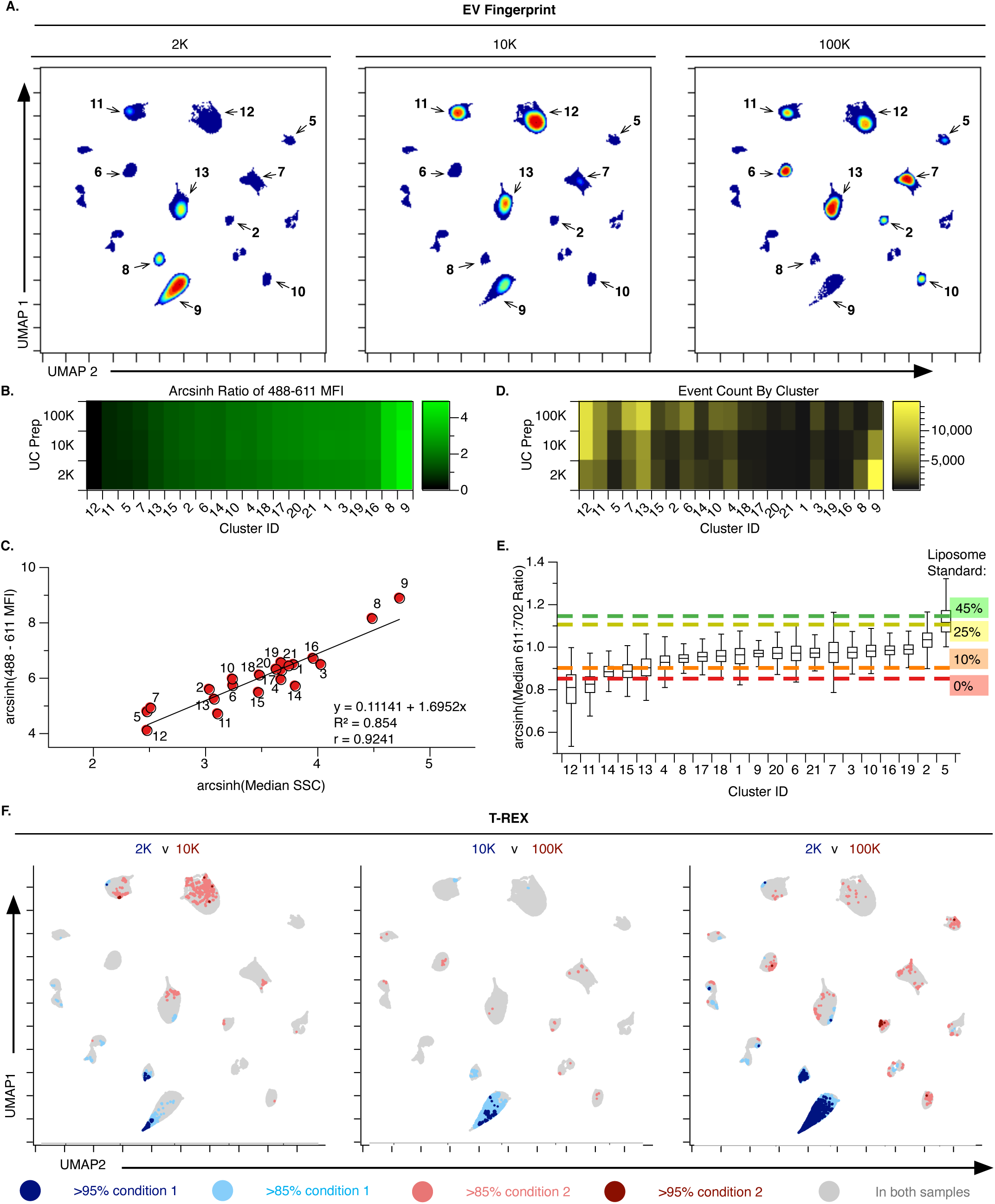
Enrichment of small and large EV populations correspond to UC preparations. **a,** EV Fingerprints of DG-UC isolated EV samples upon re-examination of 15 populations containing ≥ 1% of total EVs and removal of non-specific particles. A relative proportion of each sample (90%) was included in the re-examination (354,254 events). The EV Fingerprint revealed 21 clusters found in 2K (left), 10K (middle), and 100K (right). **b,** Heatmap of clusters separated by DG-UC speed and sorted according to 488-611 median fluorescence intensity (MFI) for each cluster. **c,** Heatmap of EV density (abundance) per cluster, using the cluster ranking from panel **b**. **d,** Cluster plot mapping 488-611 MFI to SCC (Pearson correlation coefficient r(19)=.92, p<.0001, two-tailed) **e,** Box plot visualization of spectral emission ratio (611:702 nm) for each cluster overlaid by spectral emission of liposomes with defined cholesterol content Cholesterol liposome standards (from Extended Data Fig. 3) with 0% = red, 10% = orange, 25% = yellow, and 45% = green. **f,** Independent pairwise evaluation of DG-UC samples with T-REX Statistical thresholds of >85% and >95% expansion in the KNN region denoted in red or blue respectively. *k*-value = 60.

The UMAP embedding reveals EV populations as clearly resolved clusters with sample-specific changes in particle density are readily evident (Fig. 3a). A heatmap of the 21 clusters identified ranked according to their 488-611 MFI suggests that the EV populations vary greatly in size, with cluster 9 representing the brightest and therefore the largest EV population (comparing left to right, Fig. 3b). This is confirmed when plotting fluorescence (MFI) of the population against its scatter (Fig. 3c: linear regression R^2^ = 0.854). Importantly, each of the three preparations contains detectable quantities of every EV population with similar MFI (Fig. 3b). However, the relative enrichment of each EV population is very different for each preparation (Fig. 3d). Larger EVs (high MFI and high SSC, i.e. clusters 8 and 9) are enriched in 2K preparations while smaller EVs (low MFI and low SSC, i.e. clusters 5, 7, 11, 12 and 12) are enriched in 100K with the 10K preparation containing a mixture of sizes (Fig. 3d). These observations are consistent with TRPS measurements (Extended Data Fig. 7a) and published findings^44^.

The resolution of several EV populations with very similar di8 emissions at 611 nm (e.g. 5, 7, and 11; Fig. 3c) indicates that EV Fingerprinting deconvolves EVs using multiple metrics. Indeed, a comparison of the 611/702 emission ratio as shown in Extended Data Figure 3f, reveals that clusters 5, 7, and 11 have very distinct 611/702 emission ratios (0.84, 0.98, and 1.12 respectively, Fig. 3e). When mapped against cholesterol-containing liposome standards, these ratios suggest that EVs in cluster 5, 7, and 11 have unique lipid compositions corresponding to 25%, 10%, and 0% respectively. Interestingly, while S-EVs, enriched in 100K preparations range widely in this ratiometric lipid indicator (from 0.8 to 1.12), L-EVs, enriched in 2K and 10K preparations (cluster 8 and 9) seem to be positioned at the midpoint of this range (0.95, Fig. 3e). Indeed, only the S-EVs differentially enriched at 100K can be found outside the 0.93-0.98 range that contains L-EVs (Fig. 3e).

To ensure that the observed differential enrichment of EV populations was not a product of human bias, we completed an unsupervised assessment with “Tracking Responders EXpanding” (T-REX) algorithm, which detects regions of significant change within phenotypically homogeneous events in a pair-wise comparison of two conditions, such as low- or high-speed centrifugation^45^. Using the same UMAP embeddings, three pairwise sample comparisons across 2K, 10K, and 100K DG-UC with T-REX confirmed the size-based EV enrichment identified by analysis of individual HDBSCAN clusters (Fig. 3f). The L-EV clusters 8 and 9 are identified by T-REX as enriched (≥85%) in the low-speed conditions while several S-EV clusters (including 5, 11, 12 and 13) were identified as enriched in the high-speed conditions. Thus, using EV Fingerprinting as a basis for differentiating between the EV populations, both conventional cluster analysis and T-REX, reveal that S- and L-EVs are enriched differentially by DG-UC.

### EV Fingerprinting reveals that the loss of Rab27a differentially impacts select small EV populations

EV secretion through late endosomal/lysosomal compartments relies on a small GTPase, Rab27a, which controls MVB docking to the plasma membrane^4^. Thus, the knockdown (KD) of Rab27a can reduce exosome/S-EV secretion^14^. However, whether the EV populations that change in response to Rab27a inhibition are all exosomes or include small ectosomes, is unknown. We performed a comparative EV Fingerprinting analysis of well-characterized Rab27a shRNA-mediated knockdowns (KD1 and KD2) and a scrambled shRNA control (Scr) in HT1080 cells to determine if the loss of Rab27 impacted all S-EV populations uniformly (Extended Data Fig. 8a)^14,26^.

The previously published reduction of S-EV secretion upon knockdown of Rab27a was confirmed by NTA analysis of both 10K and 100K preparations, where the expected 50% reduction in EV secretion rates was observed in the 100K preparations, but not in the 10K preparations (Extended Data Fig. 8b)^14,26^. The same 10K and 100K EV preparations from Rab27a KD and Scr were subsequently analyzed by flow cytometry after di8 labeling (Method #2). Congruent with NTA data, flow cytometry EV gated event numbers were reduced for Rab27a KDs in the 100K, but not the 10K preparations (Extended Data Fig. 8c and d). EV preparations were subsequently analyzed using the EV Fingerprinting workflow. Similar to the PC3 analysis in Figure 3, EV Fingerprinting reveals differential enrichment of EV populations across the HT1080 10K and 100K EV preparations (Scr 10K vs 100K, Fig. 4a). Among these, several 100K EV populations from Rab27a KD cells appear visibly reduced in the 100K, while 10K EV populations appear unaffected (KD1 and KD2, clusters 7,16, and 19, Fig. 4a).

**Figure 4:**
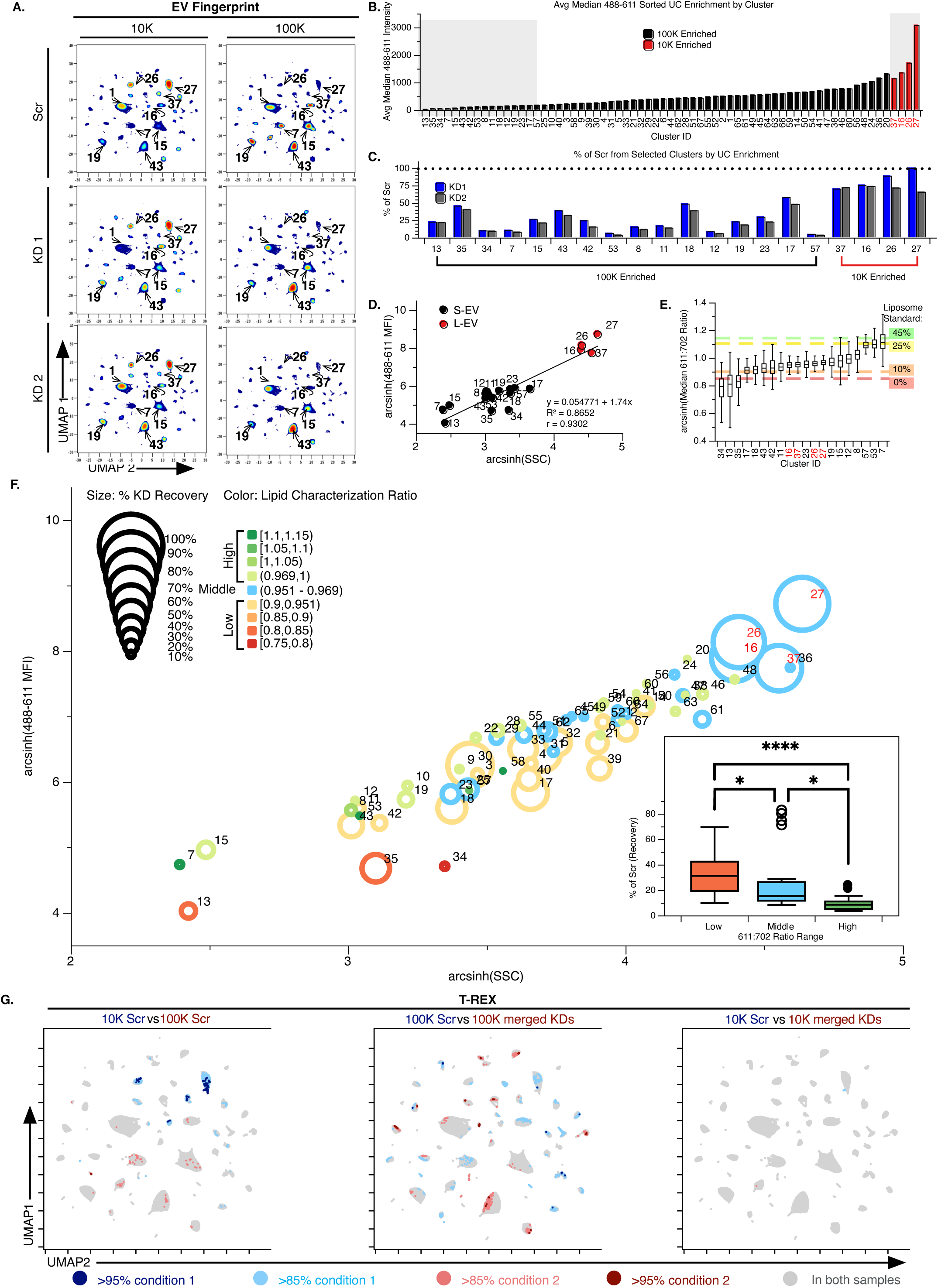
EV Fingerprinting resolves populations disrupted by the impairment of EV secretion. **a,** EV Fingerprints after re-examination of 35 selected populations using 10K (left), and 100K (right) pellets from HT1080-Scr (top), HT1080-Rab27a KD1 (middle), and HT1080-Rab27a KD2 (bottom) cells. Data were sampled relatively (80%, 1,102,755 events total) and background was removed. 67 resulting clusters were identified as shown in the plots. **b,** Cluster IDs ranked by 488 - 611 average MFI by cluster. Cluster IDs ≥50% enriched in the 100K (black) or >50% enriched in the 10K (red) defined in Extended Data Fig. 8e, correspond to S- and L-EV, respectively. Grey boxes highlight 25% of all S-EV clusters highly enriched in the 100K (left box, over black bars), or all L-EV clusters enriched in 10K (right box, over red bars). **c,** Event recovery as a percentage of Scr per cluster in KD1 (blue) and KD2 (grey) cell lines. Selected enriched clusters from L-EV (right) and S-EV (left) highlighted in (**b**) are shown. **d,** Relationship of EV 488-611 MFI and SSC by cluster. Enrichment of S-EV and L-EV is shown in black or red, respectively. Pearson correlation coefficient r(18)=.93, p<.0001, two-tailed. **e,** Lipid characterization of select clusters from (**b**) using the 611:702 ratio of emissions. Cholesterol liposome standard 611:702 ratios (from Extended Data Fig. 3) are overlaid with 0%=red, 10%=orange, 25%=yellow, and 45%=green. **f,** Comprehensive characterization of all 67 clusters identified upon Rab27a KD informed on event recovery (point size), particle size (plotted axes), and lipid composition (color). Inset categorizes low (orange), middle (blue), and high (green) lipid composition ratios with respect to % event recovery. Kruskal-Wallis test with multiple comparisons (p<0.0001) **g,** T-REX analysis on UMAP plots indicating regions of significant change in L- and S-EV enrichment (10K Scr vs 100K Scr, left), S-EV disruption (100K Scr vs 100K merged KDs, middle), and L-EV stability (10K Scr vs 10K merged KDs, right). Direction and degree of change are shown in red and blue. *k*-value = 60.

To determine if the loss of Rab27a impacted the distinct S-EV populations equally, a detailed comparison of S-EV and L-EV was pursued. For this evaluation the S-EV and L-EV populations were defined according to their relative enrichment in 100K vs 10K preparation and their 488-611 MFI as relative indicator of size (Extended Data Fig. 8e and Fig. 4b, respectively). S-EV were defined as 100K enriched EV populations in the lower quartile (least bright) of the MFI range (black, Fig. 4b). L-EV were defined as the 10K enriched EV (red, Fig. 4b). Since only four populations matched the L-EV criteria, all were included in the subsequent analyses.

The impact of Rab27a on selected (blue boxes, Fig. 4b) individual S-EV and L-EV populations were assessed by plotting KD1 and KD2 as a relative % of the control for each EV population (Scr, Fig. 4c). The overall reduction in S-EV is readily apparent, however, not all S-EV populations are impacted equally. Some EV populations retain ≤ 10% of the levels found in the control Scr cell line (i.e. 7, 53, 12, and 57, Fig. 4c) while others retain ≥ 40% of the original levels (i.e. 35, 43, 18, and 17, Fig. 4c). This variation suggests that Rab27a does not impact the release of all S-EV uniformly, which is consistent with its known role in selectively regulating the secretion of exosomes and not ectosomes.

The underlying variation of these EV populations was investigated further by exploring the correlation of EV size and lipid composition to the Rab27a-dependent EV production. Plotting the di8 488-611 MFI against SSC as a sizing metric of EVs demonstrates a strong correlation between these two metrics (R^2^= 0.8652) and validates the relative size stratification of these EV populations (Fig. 4d). Consistent with them being larger in size, L-EVs exhibit a 488-611 MFI and SSC that significantly exceeds that of S-EVs. The modest diminution of these L-EV populations (≤25% change vs. ≥50% change for S-EV populations, Fig. 4c) further confirms the observation that Rab27a KD preferentially diminishes the production of S-EVs. However, a comparison of the two largest S-EV populations (clusters 17 and 57) suggests that size is not the only factor in Rab27a-dependent production, as EV population 57 remains at 50% while population 17 is diminished to ≤ 5% of Scr output (Fig. 4c).

Since di8 fluorescence emission is lipid-dependent (Extended Data Fig. 3),^38–40^ ratiometric analysis was performed by mapping the 611:702 emission ratio from each di8-labeled EV population against a 100 nm liposome standard with increasing cholesterol content (Fig. 4e). L-EV populations (red clusters 16, 37, 26 and 27, Fig. 4e) display a 611:702 emission ratio of approximately 0.95 while the S-EV populations range widely from 0.8 to 1.1. Moreover, four of the five EV populations most impacted by Rab27a KD (a reduction of ≥90% of Scr) have a 611:702 ratio consistent with high cholesterol content (>0.95) while the EV populations least impacted by the Rab27a KD (a reduction of ≤75%) all have a 611:702 ratio consistent with low cholesterol content (<0.95) (Fig. 4e). These observations suggest that Rab27a KD preferentially diminishes the biogenesis of EVs with elevated levels of cholesterol. This is confirmed when all EV populations were mapped by size, lipid characterization ratio, and % recovery after Rab27a KD (Fig. 4f). Regardless of their size, S-EV populations with a lipid characterization ratio > 0.95 (light to dark green circles Fig. 4f) are diminished greatly when compared to S-EV with a ratio < 0.95 (orange to red circles, Fig. 4f). This is readily evident when evaluating the impact of Rab27a KD in aggregate for EVs with a high, middle, and low lipid characterization ratios (red, blue and green box plots, Fig. 4f inset). Note that in the Fig. 4f inset, the four L-EV populations (16, 26, 27 and 37) were plotted as outliers (black circles) and not included in the statical comparison of the S-EV.

Using T-REX, three pairwise comparisons were performed to assess the effects of Rab27a KD on S-EV (Fig. 4g). The pairwise comparison for L-EV preparations (10K) from scrambled vs Rab27a KD cells revealed no significant changes while a comparison for S-EV preparations (100K) identifies numerous populations impacted by the loss of Rab27a (Fig. 4f). These observations corroborate the specificity of Rab27a perturbation on S-EV while leaving the production of L-EV mostly unaffected.

### EV Fingerprinting identifies EV populations impacted by CD63 overexpression

The tetraspanin CD63 is a common EV cargo that has been shown to regulate biogenesis of S-EV^37,46,47^. To gain understanding into its role in EV biogenesis, we evaluated the impact of CD63 on vesicle heterogeneity by comparing the EV Fingerprint of HT1080 cells overexpressing CD63 (pHluorin-CD63) with that of control cells (HT1080-Parental). Consistent with Sung et. al. ^26^, we observed an increase in the numbers of EVs by NTA in the 100K EV preparation from pHluorin-CD63 cells when compared to parental cells (Parental vs pHluorin, Fig. 5a). Elevated CD63 incorporation as EV cargo from pHluorin-CD63 cells was confirmed by Western blotting (Fig. 5b). EV flow cytometry using dual labeling of di8 and anti-CD63 or control (Iso) antibody (Method #2) confirmed the increase in CD63-positive (CD63+) EVs upon CD63 overexpression from 3.57% to 33.03% of the total EV population (Parental vs pHluorin, Fig. 5c).

**Figure 5:**
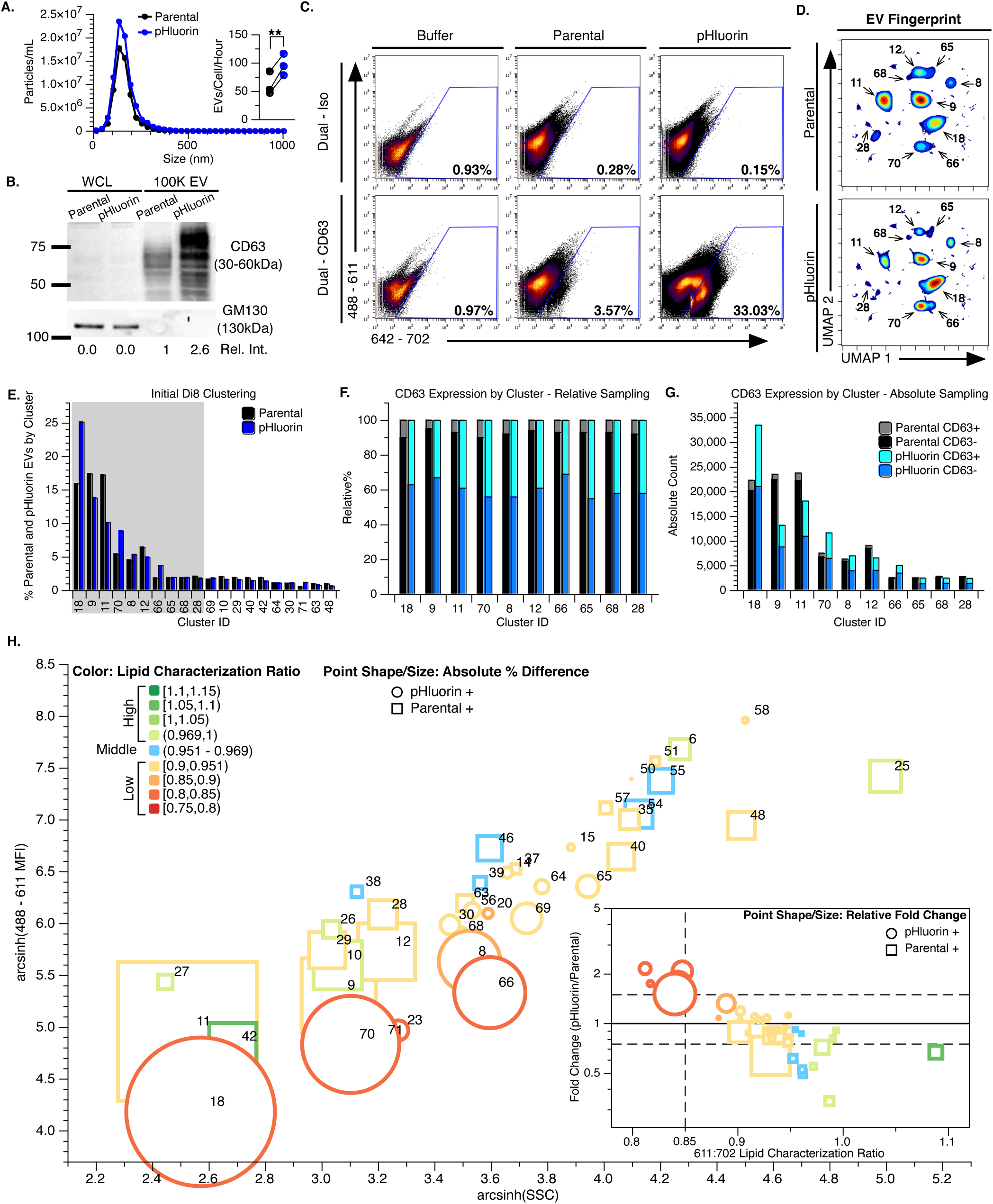
EV Fingerprinting identifies populations affected by CD63 overexpression (OE). **a,** Nanoparticle tracking analysis (NTA) showing size distribution and estimated EV secretion rate (EVs/Cell/Hour) for EVs isolated from HT1080-Parental (Parental) and HT1080 pHluorin-CD63 (pHluorin) cells. **b,** Western blotting analysis of CD63 protein expression in Parental and pHluorin whole cell lysates (WCL) and 100K EV. Relative intensity of CD63 signal in EV preps is shown below (Parental:pHluorin). WCL CD63 signal was normalized to respective GM130 expression. **c,** Representative flow cytometry scatter plots of fresh EVs and blank buffer controls dual stained with di8 (y-axis, 488-611) and antibodies (x-axis, 642-702, Isotype control (Iso) or anti-CD63 (CD63)). **d,** EV Fingerprints from initial analysis using di8 parameters (10% relative sampling, 361,483 events total) for Parental and pHluorin 100K EVs. 69 clusters were identified. Cluster IDs shown represent the top 10 populations (top 10 of ≥1% of total data) enriched according to event counts. **e,** Percentage of event counts for parental (black) and pHluorin (blue) EVs for the top 20 clusters identified in **d** (top 20 of ≥1% of total data). The top 10 clusters from **d** are highlighted in the grey box. **f-g** Individual re-examination of the top 10 clusters (shown in **d** and **e**) quantifying the relative % (**f**) and absolute number (**g**) of CD63-positive or -negative EVs for each cell type by cluster. **h,** Comprehensive characterization of all 69 clusters identified upon CD63 overexpression informed on absolute event count by cell line (point size and shape), particle size (plotted axes), and lipid composition (color). Inset stratifies low (orange), middle (blue), and high (green) lipid composition ratios with respect to fold change in pHluorin/Parental events.

EV Fingerprinting of 100K EV UC preparations from pHluorin-CD63 and parental cells readily reveals increases and decreases in distinct EV populations upon CD63 overexpression (pHluorin vs Parental, Fig. 5d). Plotting the top 20 most abundant EV populations as a portion (%) of the total illustrates that CD63 expression causes a selective increase in the relative abundance of five populations and an unexpected decrease in the relative abundance of three others (Fig. 5e). Further examination of the proportion of CD63+ vesicles (positive for both di8 and anti-CD63 antibody) in each of the top ten EV populations demonstrates that overexpression of CD63 leads to a universal 4-fold increase in the proportion of CD63+ EVs (Fig. 5f). However, while the proportion of CD63+ EVs increased relatively uniformly across all EV populations (Fig. 5f), the absolute number of EVs in each population did not (Fig. 5g). Of the top 10 EV populations, only four are increased in abundance upon expression of CD63 (18, 70, 8 and 66) while three are decreased in abundance (9,11, and 12).

Ratiometric 611:702 analysis for all clusters was performed to determine if the selective impact of CD63 corresponded with EV lipid composition. Changes in EV biogenesis were visualized in Figure 5h by mapping EV clusters by particle size (488-611 MFI vs SSC), lipid characterization ratio (611:702 ratio), and the % increase/decrease (as a proportion of total EV production) upon CD63 overexpression. In this visualization, it is evident that the largest changes in EV biogenesis occur for smaller EVs (lower left quadrant, Fig. 5h). However, increases in biogenesis are only seen in EV populations with a lipid characterization ratios <0.85 and decreases in EV biogenesis are only observed in EV populations with a ratio >0.85 (Fig. 5h inset). These observations reveal that the increase in EV biogenesis upon CD63 expression is selective as it occurs primarily for the smaller EVs with a lipid characterization ratio consistent with low cholesterol content (Red circles, Fig. 5h inset).

### Detection of cancer-associated L-EV populations in human plasma using EV Fingerprinting

Circulating EVs present in plasma are thought to be a systemic reflection of health and disease and thus a promising target for liquid biopsies^48–50^. However, the abundance of other components in plasma, including proteins and lipoproteins, makes the analysis of plasma EVs especially challenging ^51^. To determine if EV Fingerprinting could deconvolve a complex sample and identify tumor-derived EVs in plasma, we applied this method to human plasma spiked with L-EVs derived from PC3 cells. PC3 cells are known to generate a L-EV population, termed LO, associated with aggressive disease^5,37,52–54^. LO-containing EV preparations were isolated from PC3 cells using 10K d-UC and characterized using TRPS (Extended Data Fig. 9a). The ability to detect PC3-derived LO in plasma was subsequently assessed by EV flow cytometry (Fig. 6 and Extended Data Fig. 9b). For the purpose of tracking L-EV, they were defined as particles with a 488-611 MFI greater than that of the 400 nm liposome standard (Fig. 2d,e and Extended Data Fig. 3b,c). Using this threshold, the addition of L-EV from 10K PC3 EV preparation to plasma was readily identified and quantified (Fig. 6a,b). EV Fingerprinting was subsequently used to deconvolve EV populations present in plasma before and after the addition of PC3-derived EVs. As in Fig. 3, EV Fingerprinting of PC3 revealed a prominent L-EV population (Cluster 13, Fig. 6c,d,e) composed of the largest EVs produced by PC3 and thus consistent with previously described LO^5,37,54^. The addition of these tumor-cell derived EVs to plasma is readily detected (Fig. 6d). Corroborating this observation, the unbiased analysis of the plasma EV Fingerprint with and without PC3-derived L-EVs using T-REX readily identifies the increase of EVs in cluster 13 (Fig. 6f).

**Figure 6:**
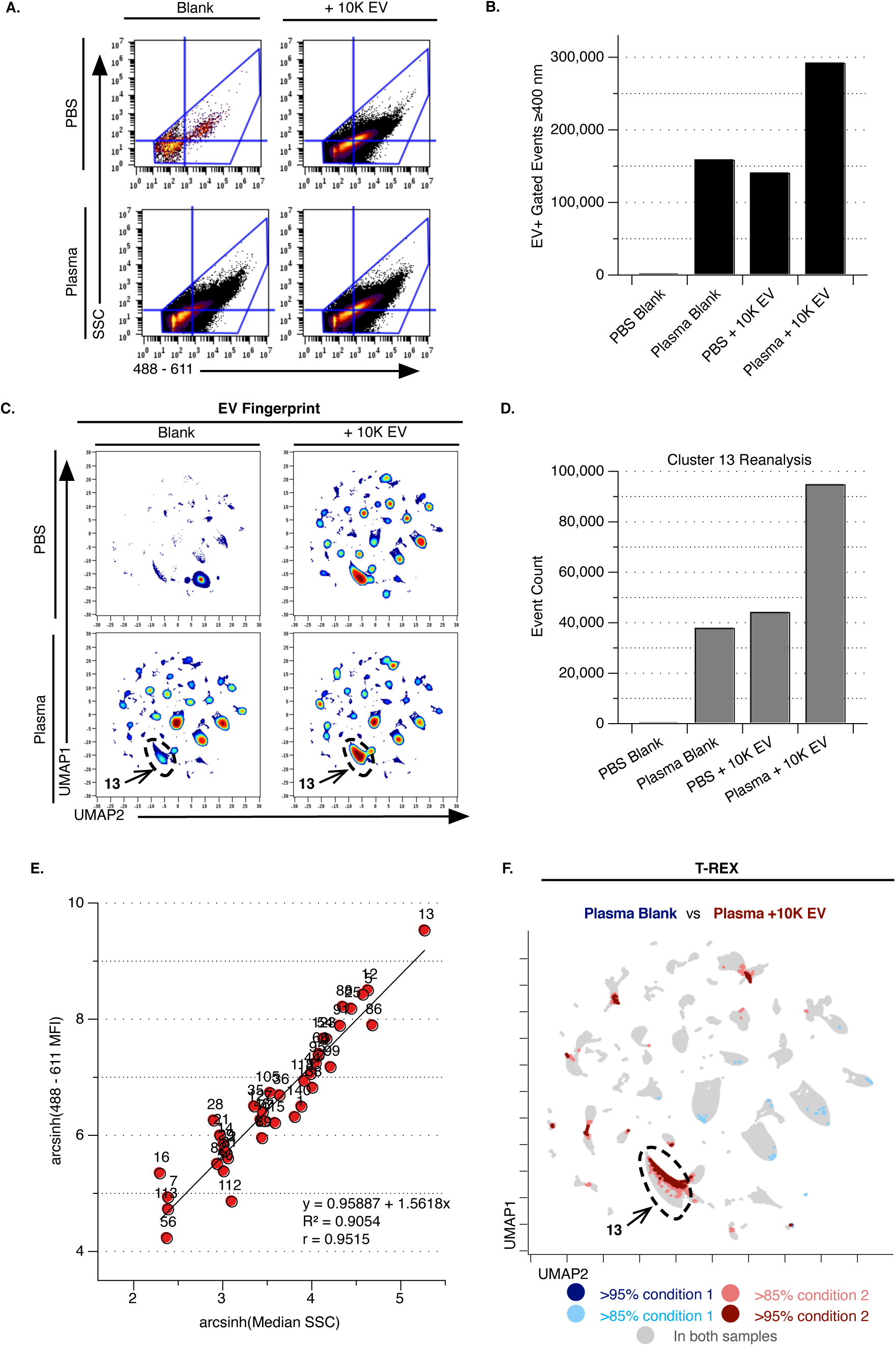
Quantitative detection of tumor cell-derived Large EVs in plasma through EV Fingerprinting and T-REX. **a,** Conventional flow cytometry gating of L-EVs (d-UC isolated 10K PC3 EVs) spiked into plasma (+10K EVs). EV+ event gating (polygon gate) to quantify L-EV+ events (upper right quadrant gate) as described in Extended Data Figure 9b. Data were collected for 10 min using a modified Method #2 di8 staining of EVs freeze/thawed at −80℃. **b,** Quantitation of EV+ L-EV+ events from (**a**). **c,** EV Fingerprints of initial analysis (absolute sampling, 200,000 events per samples for a total of, 610,123 events) of PBS blank, PBS + 10K EV, plasma blank, and plasma + 10K EV. **d,** Quantitation of events upon re-examination of Cluster 13 alone using di8 features and 80% relative sampling from each condition (225,263 events total). **e,** Size characterization of EV populations. Pearson correlation coefficient r(37)=.95, p<.0001, two-tailed. **f,** UMAP from (**c**) with T-REX analysis of “Plasma blank” and “Plasma + 10K EV” samples. Enrichment is shown in blue and red. *k*-value = 60.

## Discussion

Single-EV analysis is a critical emerging strategy that is needed to resolve individual EV populations and define their association with specific biology. We developed EV Fingerprinting as a method of high-throughput, single-EV analysis that enables deconvolution of EV populations and their characterization from complex biological samples. Like other flow cytometry methods, EV Fingerprinting distinguishes itself from bulk EV analysis methods by allowing for the sizing and cargo characterization of individual EVs. However, unlike other flow methods, EV Fingerprinting leverages dimensional reduction of 20 parameters collected from the fluorescent lipophilic dye, di8, to cluster EVs into populations based on shared parameters, beyond the single metric of size. The result is a quantifiable stratification of EV populations informed by lipid composition that enables the discovery of population-specific changes in response to molecular perturbation. This is a critical technological advance that enables detailed EV analysis without prior knowledge of the target EV population or the need for purification prior to analysis.

We demonstrated the utility of EV Fingerprinting by revealing that specific molecular perturbation of EV biogenesis impacts select EV populations (Fig. 4 and 5). Moreover, using ratiometric analysis of emission at 611 and 702 nm, we were able to demonstrate that Rab27a KD preferentially decreased the biogenesis of EVs with a lipid characterization ratio consistent with high cholesterol content. Conversely, CD63 expression preferentially increased the biogenesis of EVs with a ratio consistent with low cholesterol content. The authors observed that the role of CD63 in EV biogenesis is more nuanced than simply being EV cargo because elevated CD63 expression alters the biogenesis of select EV populations, but it is universally incorporated as cargo into all EV populations identified. EV Fingerprinting of EVs purified by DG-UC quantitatively stratifies the selective enrichment of EVs by size and demonstrates the broad heterogeneity of tumor-derived EVs. Subsequent analysis of human plasma spiked with DG-UC enriched tumor cell-derived EVs demonstrates the capacity of EV Fingerprinting to resolve EV populations in complex biological samples and detect tumor-derived EV.

EV Fingerprinting still has some limitations and potential for evolution. Data acquisition relies on single particle detection by fluorescence triggering and is thus limited by the instrument’s ability to discern the particle from background. For samples with high background, we recommend dilution of the sample or removal of contaminants to improve sensitivity. One example of this is lipoproteins in plasma that are highly abundant and incorporate di8 dye. The two-phase Fingerprinting workflow does enable the digital exclusion of background particles in the downstream analysis, however, non-specific particles extend instrument runtime and sometimes require additional dilution in order to capture sufficient data for specific EVs. Thus, EV enrichment methods such as d-UC, DG-UC, or size exclusion chromatography may be needed. In future implementations, increased throughput on the instrument or more EV-specific staining methods may diminish the impact of non-specific particles.

The high resolution of EV Fingerprinting permits the identification of EV populations associated with specific biological functions and/or perturbations. While further standardization will improve our understanding of lipid-based clustering, EV Fingerprinting serves as a central tool to uncover the biological foundations of physiological and pathological EV heterogeneity. We envision significant potential for this methodology in studying basic EV lipid composition and biogenesis, translational research such as viral infection and therapeutic response, as well as clinical research for use as a non-invasive liquid biopsy.

## Methods

### Cell culture and reagents

HT1080 fibrosarcoma cells were maintained in Dulbecco’s Modified Eagle Medium (DMEM) high glucose (10-013-CV, Corning) supplemented with 10% bovine growth serum (BGS, SH30541.03, HyClone). HT1080 cells carrying the scrambled control or Rab27a-specific shRNAs (KD1 and KD2)^14^ or expressing pHluorin_M153R-CD63^26^ were cultured under the same conditions as the parental line. The PC3 cell line was obtained from the American Type Culture Collection (ATCC) and cultured in DMEM (Invitrogen). The DMEM was supplemented with 10% fetal bovine serum (Denville Scientific), 2 mM L-glutamine (Invitrogen) and 1% PenStrep (Invitrogen). The cells were grown at 37°C and 5% CO2. Cell viability of the EV-producer cells was tested with the 0.4% Trypan Blue (Sigma) exclusion method. All cell lines were routinely tested for mycoplasma contamination by using the MycoAlert PLUS Mycoplasma Detection Kit (Lonza).

Antibodies utilized included: anti-Rab27a (69298, Cell Signaling, 1:1,000 for WB), anti-β-actin (Ac-74, Sigma-Aldrich, 1:5,000 for WB), anti-CD63 (ab68418, Abcam, 1:500 for WB), anti-GM130 (610822, BD Biosciences, 1:250 for WB). Horseradish peroxidase (HRP)-conjugated goat anti-mouse IgG (W4021, 1:10,000 for WB) or goat-anti-rabbit IgG (W4011, 1:10,000 for WB) were purchased from Promega. APC anti-human CD63 (353008, 1µg/mL for FC) APC mouse IgG1, k isotype Control (400120, 1µg/mL for FC) were purchased from Biolegend.

### Extracellular vesicle isolation from cultured cells

Differential ultracentrifugation (d-UC) isolation was performed on HT1080-derived EV preparations as previously reported.^14,26^ Cell culture media was collected from cultures maintained at 80% confluent cells for 48 h in Opti-MEM (31985070, Thermo Fisher Scientific). The conditioned media were centrifuged at 300 × *g* for 5 min and 2,000 × *g* for 20 min to sediment live cells and cellular debris, respectively. The supernatant was centrifuged at 10,000 × *g* for 30 min (Ti45 rotor, Beckman Coulter) to collect large EV or 10K pellets. The supernatant from 10,000 × *g* centrifugation was further centrifuged at 100,000 × *g* for 18 h (Ti45 rotor) to pellet small EV or 100K pellets. Both 10K and 100K pellets were resuspended in phosphate-buffered saline (PBS) and spun again at 10,000 × *g* for 30 min or 100,000 × *g* for 3 - 6 h.

The isolation and density gradient purification of PC3 EV was performed as reported with minor modifications described below.^5,44,55,56^ PC3 cells were grown in 18 × 150 mm cell culture dishes (Corning) until 90% confluence, washed in PBS and serum-starved for 24 hours before the collection of cell conditioned media in serum free-medium (same DMEM as cell culture conditions, minus serum). The conditioned media was centrifuged at 300 g x 3, 5 min each, to pellet down floating cells, followed by centrifugation at 2,800 g for 10 min to pellet 2K large EV. The resulting conditioned media was spun in an ultracentrifuge at 10,000 g for 30 min (k-factor 2547.2) for the collection of 10K large EV and the supernatant was then spun at 100,000 g for 60 min (k-factor 254.7) for the collection of 100K small EV. All d-UC steps were performed at 4°C. 2K, 10K, and 100K pellets were either resuspended in 0.2 µm-filtered PBS and used as d-UC pellets or subjected to further Optiprep™ (Sigma) density gradient purification. Fresh pelleted EV were resuspended in 0.2 µm-filtered PBS and deposited at the bottom of an ultracentrifuge tube. Next, 30% (4.3 mL, 1.20 g/mL), 25% (3 mL, 1.15 g/mL), 15% (2.5 mL, 1.10 g/mL), and 5% (6 mL, 1.08 g/mL) iodixanol solutions were sequentially layered at decreasing density to form a discontinuous gradient. Separation was performed by ultracentrifugation at 100,000 g for 3 h 50 min (4°C, k-factor 254.7) and EV-enriched fractions collected either at 1.10–1.15 g/mL for large EV or 1.10 g/mL for small EV.^5^ Purified EV were then washed in PBS (100,000 g, 60 min, 4°C) and resuspended in 0.2 µm-filtered PBS. All ultracentrifugation spins were performed in a SW28 swinging rotor (Beckman Coulter).

### Extracellular vesicle characterization

HT1080-derived pellets were resuspended in PBS and used fresh for NTA using ZetaView (Particle Metrix), Western blotting, or flow cytometry using the Amnis CellStream™ (Luminex). EV-specific marker expressions were validated in previous studies from our group.^14,26^ PC3-derived pellets were resuspended in PBS and characterized fresh through TRPS using qNano (Izon) and subsequently frozen at −80 ℃ before flow cytometry analysis.

### Liposome Preparation

Liposomes were prepared through the established extrusion method.^57^ Briefly, lipids were dessicated for 2 hours and allowed to reach RT. 1,2-distearoyl-sn-glycero-3-phosphocholine (DSPC), Cholesterol and 1,2-Dimyristoyl-rac-glycero-3-methoxypolyethylene glycol-2000) (DMG-PEG2000) were weighed at the indicated ratios (Supplementary Tables 4,5) and dissolved in chloroform, evaporated using a nitrogen stream and left under vacuum overnight to form a lipid film. This film was rehydrated with PBS (pH 7.4) at 65 °C for 2 hours with vortexing (every ~15 mins). These multilamellar vesicles were further processed with 5 freeze (liquid nitrogen) and thaw (65 °C water bath) cycles. The nanoparticles were extruded through stacked polycarbonate filters (400, 200 or 100 nm) at least 10 times. The size of liposomes was measured using dynamic light scattering (Malvern Zetasizer).

### EV Staining for Flow Cytometry

#### Di-8-ANEPPS Preparation and Storage

Lyophilized di8 powder (Biotium, Cat#: 61012) was resuspended in DMSO (Millipore Sigma, Cat#: D8418) to a concentration of 5000 µM. The dye was then filtered with a 0.2 µm regenerated cellulose syringe filter (Corning, Cat#: 431215) and stored at −20 °C in 25 µL aliquots. Only the aliquot actively being used was stored short term at 4 °C. This 5000 µM stock was diluted fresh in DMSO before each experiment to make the 25 µM working stocks used for staining buffers.

#### Molecular Crowding (MC) Buffer Preparation and Storage

A 6.5% w/v solution of dextran, mr= ~100,000 (Millipore Sigma, Cat#: 09184) was prepared in DPBS without calcium and magnesium (Corning, Cat#: 21-031-CM) to make 2x MC Buffer and filtered with a 0.2 µm PVDF bottle filter (Millipore Sigma, Cat#: S2GVU05RE) in a sterile hood. 50 mL aliquots were stored at 4 °C until use.

### Staining Strategies

The staining strategy used was dependent upon the approximate particle concentration of the sample. The methods are outlined below.

Method #1 (Extended Data Fig. 1a): This staining method was used for samples with a low particle concentration (e.g. conditioned media or dilute purified EV) and is not amenable to multiplexing with antibody stains. EV-containing samples and serial dilutions were prepared in MC buffer and combined with the staining buffer prior to analysis (See Protocol). The final staining volume for each sample was 100 µL (50 µL of sample + 50 µL of staining buffer) and final di8 staining concentration was 0.25 µM. First, the staining buffer was prepared by diluting 25 µM di8 working stock into a 2x MC buffer for a dye concentration of 2.5 µM (1:50 in 50 µL of 2x MC per sample). Staining buffer was spun at 16,100 x *g* (max speed) at RT for 15 min. In the meantime, sample dilutions were prepared in PBS for a final volume of 50 µL. Finally, 50 µL spun 2x staining MC buffer was added to 50 µL of diluted sample for a final staining volume of 100 µL, and di8 concentration of 0.25 µM. Samples were stained at RT in the dark for 1 hour, then directly distributed in a 96-well round bottom plate (Fisher Scientific, Cat#12565500) for flow analysis.

Method #2 (Extended Data Fig. 1b): This staining method was developed for samples with a high particle concentration (ie. Purified EV samples, most synthetically derived nanoparticles, and liposomes) and supports the use of antibody staining. In principle, staining buffer and samples dilutions were prepared separately and combined for staining as in Method #1, but also includes a post-stain 1:200 dilution. The final staining volume for each sample was 25 µL (14.5 µL staining buffer + 10.5 µL diluted sample OR 14.5 µL staining buffer + 8 µL diluted sample + 2.5 µL antibody) and staining concentration of di8 was 2 µM. First, the staining buffer was prepared by diluting 25 µM di8 working stock into 2x MC buffer. If used, antibodies were diluted in PBS to specified concentrations. Staining buffer and diluted antibodies were spun at 16,100 x *g* (max speed) at RT for 15 minutes. In the meantime, sample dilutions were prepared in PBS. Diluted sample, spun staining buffer, and spun antibody (if used) were then combined for the final di8 concentration of 2 µM, vortexed briefly and quick spun to make sure the full 25 µL staining volume was collected at the bottom of the microcentrifuge tube, and stained for 1 hour (-antibodies) or 3 hours (+antibodies) at 37 °C. Post-stain dilution tubes were then prepared with 1 mL of 1:1 x MC buffer/PBS solution. Once staining was complete, samples were diluted 1:200 using the prepared post-stain dilution tubes. Samples were then distributed in a 96-well round bottom plate for flow analysis.

#### NP-40 EV Lysis

EVs were stained using Method #2 above. After staining 1 hour at 37 °C (prior to post-stain dilution of 200x), 5 µL of 3% NP-40 was added to the stained sample making a final lysis volume of 30 µL at 0.5% NP-40. Samples were lysed for 15 minutes at RT, vortexing every 5 min. Finally, post-stain dilution of 200x proceeded as in staining Method #2.

### Flow Cytometry Data Acquisition using the Amnis CellStream**™**

The Amnis CellStream™ was calibrated and initialized before each run according to the manufacturer’s instructions. Acquisition settings were then configured as follows: Small Particle Mode = ON, Flow Rate = SLOW (3.66 µL/min), Thresholds = ZERO, Trigger Channels = NONE, FSC/SSC Laser Power = 1%, 488 Laser Power = 25%, and if antibodies were used 642 Laser Power = 50%. Stopping criteria was set to TIME and data was collected for specified durations by experiment. Compensation was not performed.

### Data Analysis

Initial data review and quality control was performed in Cytobank.^58^ Samples were assessed for event count and signal intensity across dilutions. Specific signal was delineated from background by comparing di8 and antibody-stained samples with buffer, unstained EV, and isotype controls. Data were plotted upon arcsinh transformation. Samples with event counts in the quantitative range (Extended Data Fig. 6) were selected for analysis in the EV Fingerprinting workflow and subsequent data output was uploaded to Cytobank for visualization (Extended Data Fig. 4).

#### EV Fingerprinting analysis workflow

The analysis workflow was executed in the KNIME Analytics Platform^41,42^ using Python 3.6 running on a Mac Pro (early 2008, with 14 GB 800 mHz DDR2 FB-DIMM memory connected to an external storage array (RAID5).

The workflow consists of two stages (Extended Data Fig. 4). In the first stage a randomized portion of each .fcs files is processed for exploration. In the second stage, clusters of interest can be re-examined by selecting and filtering clusters from the full data set for comprehensive analysis. In each stage the flow data is arcsinh transformed and embedded in 2D (UMAP)^27,28^, followed by cluster identification (HDBSCAN)^29,30^. Both UMAP and HDBSCAN algorithms can be parameterized externally (Supplementary Table 6). Re-examination was executed similarly, using XGBoost to filter EVs belonging to the clusters of interest from the original sample files. For experiments with antibody staining, the antibody emission parameters were included in the parameter selection, in addition to the di8 parameters (Supplementary Table 2). EV population classification results were exported as .csv files for downstream analysis.

#### EV population analysis

Individual data files for each analyzed sample as well as a table containing data grouped by Cluster_ID were written and exported as .csv files. UMAP plots were graphed in Cytobank by re-uploading individual data files using the Cluster_ID’s as “automatic cluster gates,” a column recognized by Cytobank as designated clusters for gating.

#### Statistical analyses

Statistical analyses and other graphing were performed in Excel (version 16.60), Prism (version 9.2.0), R (version 4.1.3) and DataGraph (version 4.7.1). 488-611 MFI by cluster was calculated from the grouped data table by including all samples in the cluster, excluding buffer controls. We considered an EV positive for an antibody when the number of events were absent or low (<1% of total) compared to the negative controls (dual stained buffer + antibody, dual stained EV sample + antibody). Statistical assessment of one biological replicate (n=1) are shown in main figures. Qualitative assessment of trends was performed on biological replicates (n ≥ 2) for each experimental condition. In total, at least three biological replicates were analyzed for each experiment (n ≥ 3). Technical replicates were included in all experiments as serial dilutions.

#### T-REX analysis

An independent pair-wise analysis of changes in EV populations was performed using a newly developed machine learning algorithm: T-REX (Tracking Responders EXpanding)^45^ in R (version 4.1.3). The data input for T-REX was a pair of samples in .csv format generated by the EV Fingerprinting pipeline for each comparison of interest. In a comparison, T-REX equally sampled events from the paired data and used KNN with a k-value of 60 to find regions of difference between samples on the UMAP axes. These “hotspots” reflect both positive regions (elevation under condition one) and negative regions (elevation under condition two) across the pairwise sample comparisons. For KNN regions containing greater than or equal to 95% of events from one of the samples (dark blue for sample 1 and dark red for sample 2), events were clustered using DBSCAN (eps = 1, minPts = 1) ^59^.

### Degree of MISEV compliance

Preanalytical variables such as isolation conditions^5,14,26,44,55,56^ and EV storage prior to analysis were reported as recommended by MISEV guidelines except for cell viability and number, which was not quantified but monitored visually^23–25^. Size-based nomenclature was used unless morphology and protein cargo indicated specific cellular origination (e.g., Golgi). Protein markers used in western blot conform to MISEV recommendations. Limitations to optical imaging and other EV characterization methods were reported. Sample preparation and staining, including assay controls, were detailed in the methods section. EV detection was assessed using serial dilutions to determine the quantitative range for every experiment and detergent-treated EV samples. The CellStream™ instrument was calibrated prior to every experiment using vendor beads. Instrument settings for data acquisition were reported for every experiment. MESF units of fluorescence were not reported due to the spectral nature of di-8-ANEPPS. Size calibration was not performed, instead fluorescence vs SSC was used as a relative measure of EV size by cluster. EV characterization was performed using the EV Fingerprinting methodology; observations made by dimensional reduction of 20 parameters were further characterized by mapping defined EV populations against sizing and lipid composition standards.

### Code availability

All codes related to EV Fingerprinting can be found at [github repository will be provided at time of publication].

## Acknowledgements

This work was supported in part by grants from the National Institutes of Health (R01CA218526 to AZ and DDV, R01CA234557 to DDV, R01CA249424, R01CA249684, U01CA224276, P01CA229123 to AMW) and the National Science Foundation (MCB-2036809 to AMW and JTW).

---

**Extended Data Figure 1:**
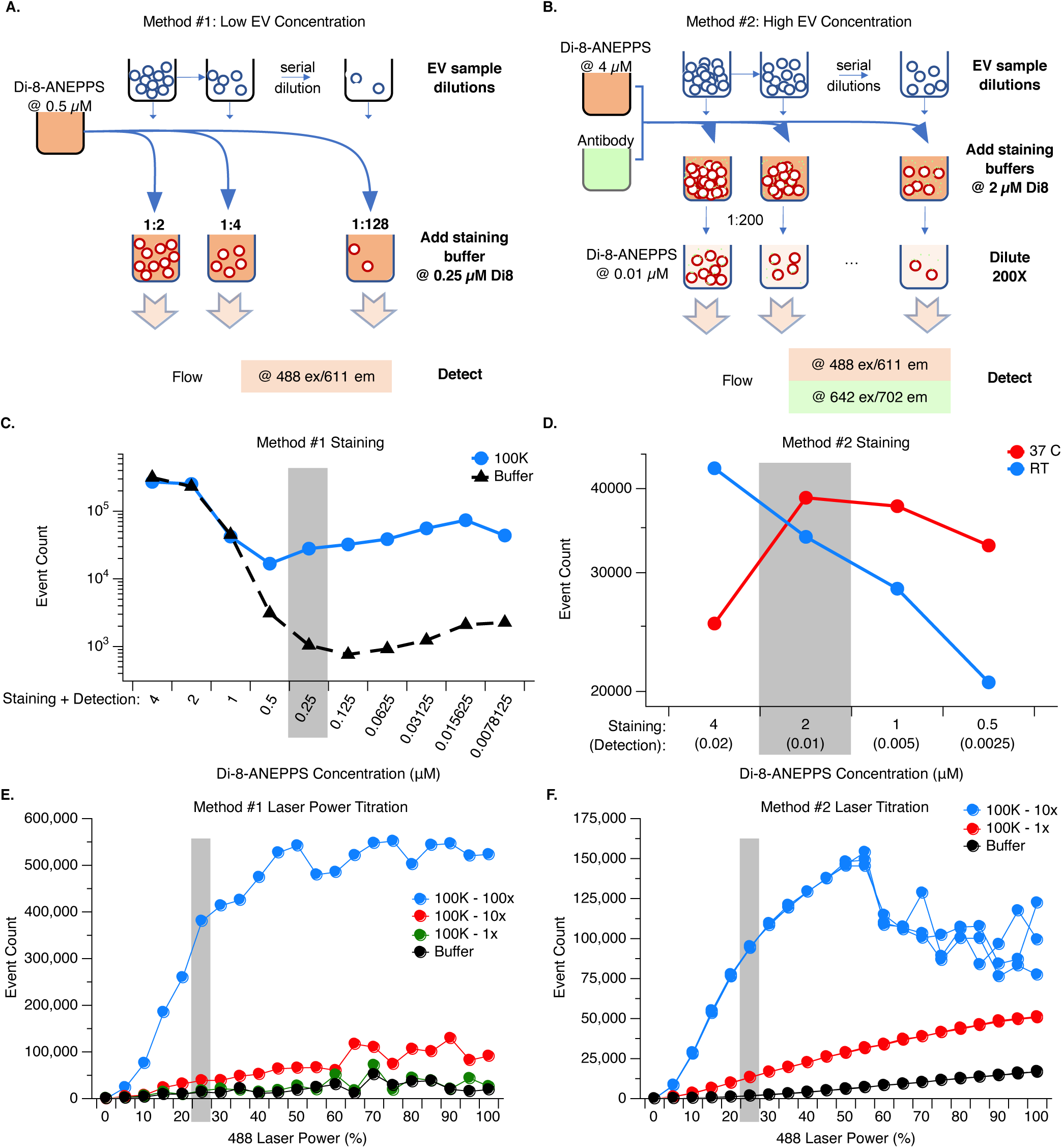
Optimized assay conditions for detecting EV by flow cytometry with Di-8-ANEPPS. **a, b,** Schematic of di8 staining of EVs using Method #1 (**a**) and Method #2 (**b**) staining strategies. Method #1 is employed for EV samples of low concentration and does not include a post stain dilution, therefore di8 staining concentration is the same during sample detection. Method #2 is used for concentrated EV samples and includes a 200x post stain dilution. **c,** Method #1 staining of a single HT1080 100K EV sample (blue circles) and buffer control (black triangles) across a 2-fold serial titration of di8 final concentrations. Grey box indicates selected staining concentration of di8 (25 µM). Raw event counts are plotted for each sample. Data were collected for 2 minutes. **d,** Method #2 staining of HT1080 100K EVs at 37°C (red) and RT (blue) upon 2-fold serial titration of di8. Grey box indicates selected di8 staining concentration and 200x diluted detection concentration (2 µM and 0.01 µM respectively). Gated event counts are shown to exclude nonspecific background from the buffer for each respective staining condition. Data were collected for 2 min. **e,** Laser power titration of Method #1 stained (0.25 µM di8) HT1080 100K EV sample. Three dilutions (blue, red, and dark green) of the same 100K EV sample were acquired at each 488-laser power including a stained buffer control (black). SSC and FSC laser settings (laser power = 1%) remained constant as well as trigger parameters (Trigger = ALL). Grey box highlights selected 488-laser power (25%). Raw event counts are plotted for each sample. Data were collected for 2 min. **f,** Laser power titration of Method #2 stained (2 µM di8 stained then diluted 200x for detection at 0.01 µM di8) HT1080 100K EV sample. Two dilutions (blue and red) of the same 100K EV sample were acquired at each 488-laser power as well as a stained buffer control (black). SSC, FSC, and trigger parameters remained constant (1%). Grey box highlights optimal 488-laser power (25%). Raw event counts are plotted for each sample. Data were collected for 2 min.

**Extended Data Figure 2:**
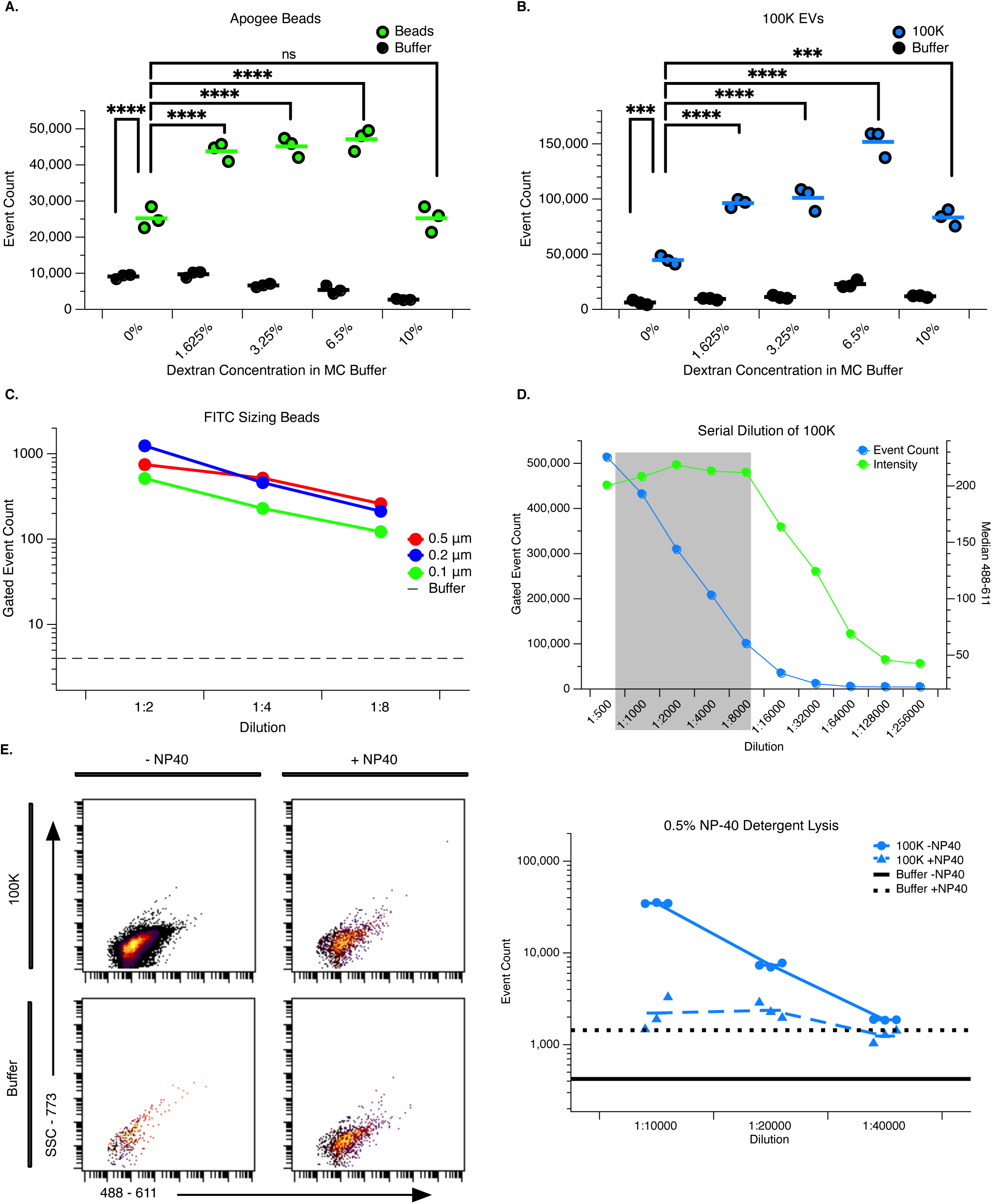
Validation of quantitative particle detection using flow cytometry. **a,** Detection of Apogee Beads (mixture of sizes, some FITC labeled, (green)) and molecular crowding (MC) buffer control (black) in different dextran staining concentrations. Raw event counts are plotted for each sample type. Data were collected for 2 min and 488-laser power was set to 25%. 2-way ANOVA with multiple comparisons, **** = P<0.0001 **b,** Detection of a HT1080 100K EV sample stained using Method #2 and MC buffer control (black) in different dextran staining concentrations. Raw event counts are plotted for each sample type. Data were collected for 2 min and 488-laser power was set to 25%. 2-way ANOVA with multiple comparisons, *** = P=0.0003 **** = P<0.0001 **c,** Quantitative detection of individual FITC-labeled sizing beads (red, blue, and green solid) across serial dilutions and buffer control (black dotted). Beads were diluted in MC buffer for a final concentration of dextran of 3.25%. Gated event counts are shown. Data were collected for 2 min at a 488-laser power of 50%. **d,** Quantitative detection range of Method #2 stained HT1080 100K EV. Gated EV+ Event count (blue) is plotted on the left y-axis and 488 median fluorescence intensity (MFI, green) is plotted on the right y-axis. Data were collected for 2 minutes using EV Fingerprinting acquisition settings (Small Particle Mode, 488-laser power = 25%, FSC/SSC= 1%, Trigger = All). The grey box indicates the quantitative detection range where MFI is stable across dilutions and event count decreases according to the dilution factor. **e,** Loss of EV detection upon lysis with 0.5% NP-40. Representative flow cytometry scatter plots (left) of gated Method #2 stained HT1080 100K EVs (blue) and buffer (black) with (+) and without (−) NP-40. Quantitation (right) of gated events across serial dilutions using three technical replicates. Data were collected for 1 min.

**Extended Data Figure 3:**
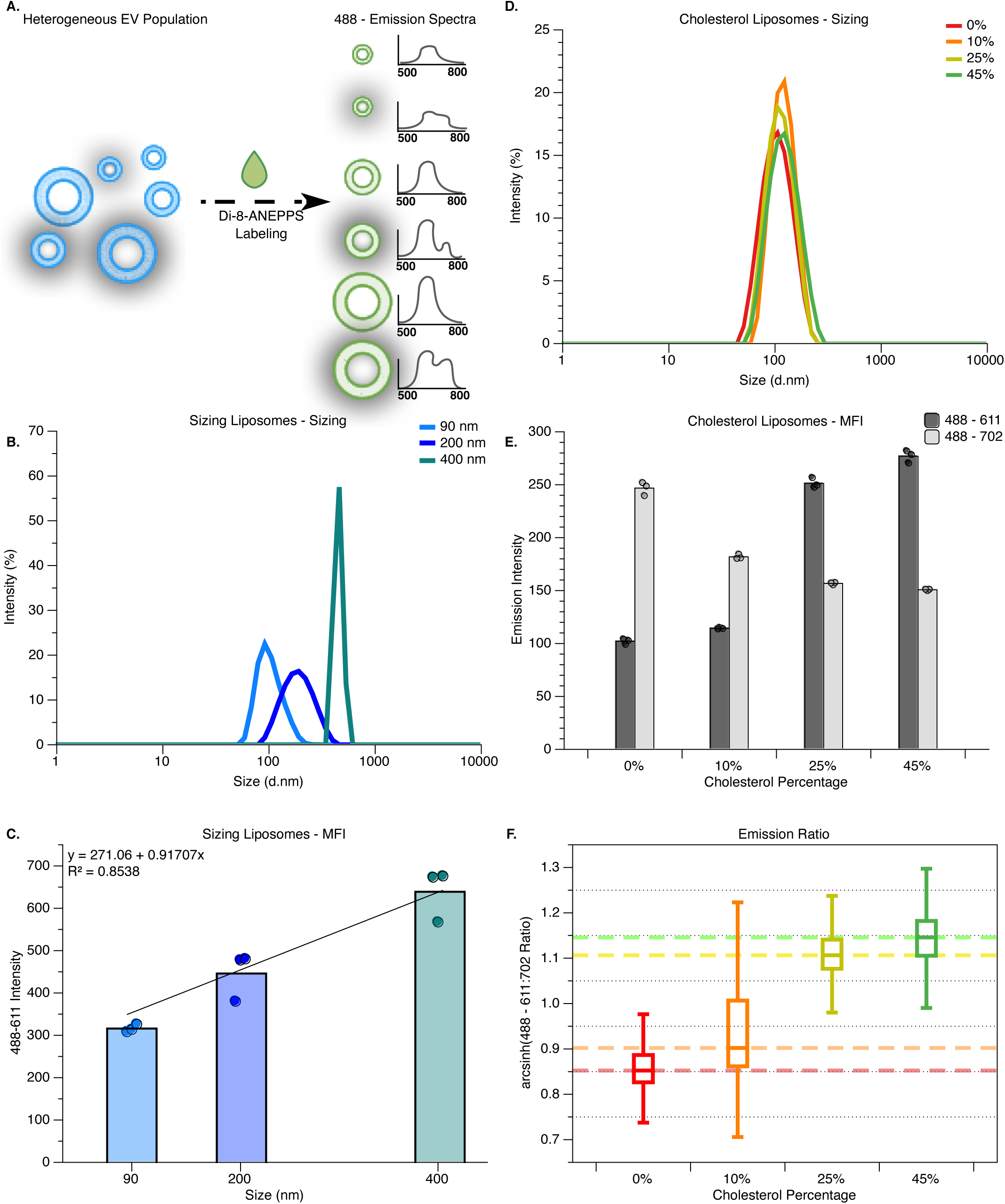
Spectral emissions of Di-8-ANEPPS reflect particle size and lipid composition. **a,** Schematic representation of heterogeneous EV labeling and the resulting emission output. The effects of EV size and lipid composition are graphically represented in collected data. **b,** Dynamic light scattering (DLS) showing size distribution of sizing liposomes (90 nm, 200 nm, and 400 nm) made of the same lipid composition. **c,** Fluorescence characterization of sizing liposomes from (**b**) using 488-611 median fluorescence intensity (MFI) determined by flow cytometry. Data were collected for 5 min using EV Fingerprinting acquisition settings. **d,** DLS sizing determination of cholesterol liposomes made of different lipid compositions (0%, 10%, 25%, and 45% cholesterol, all 100 nm). **e,** Fluorescence characterization of cholesterol liposomes from (**d**) using 488-611 MFI and 488-702 MFI determined by flow cytometry. Data were collected for 5 min using EV Fingerprinting acquisition settings. **f,** Emission ratios of cholesterol liposomes from (**d** and **e**). The 611:702 emission ratios were calculated upon arcsinh transformation of MFIs.

**Extended Data Figure 4:**
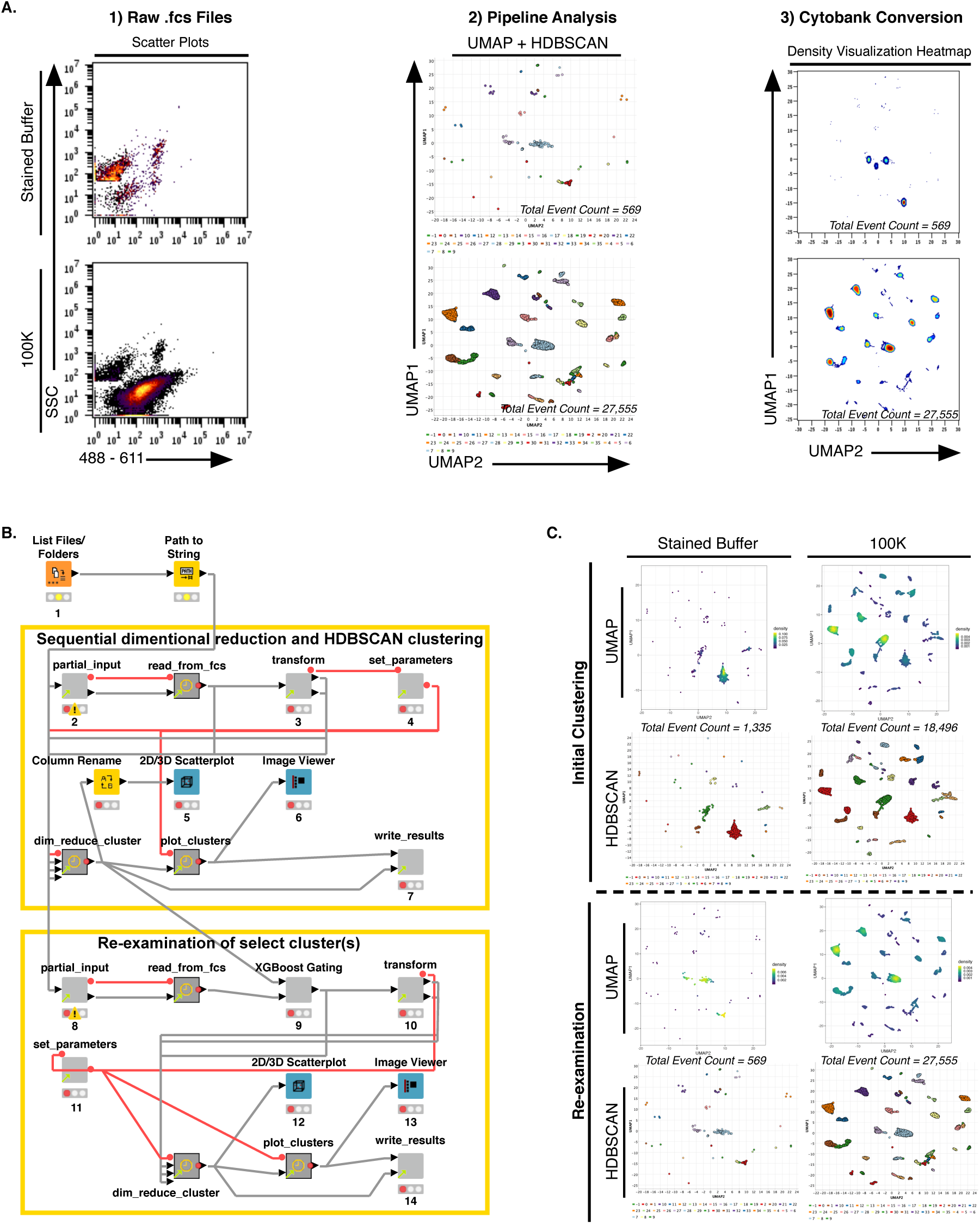
Semi-automated analysis pipeline for dimensional reduction and clustering of EV populations. **a,** Workflow of data analysis and processing after collection. Raw flow cytometry data (left, .fsc files) are uploaded and analyzed using the EV Fingerprinting pipeline in KNIME. Data are reduced in dimension (UMAP) and cluster identified (HDBSCAN) then output is exported as .csv files (middle). The .csv files are uploaded to cytobank for final visualization (right). **b,** The EV Fingerprinting pipeline in KNIME. Each node performs a data processing step that is sequentially executed to generate results. The pipeline is split into two parts: Initial clustering (“Sequential dimensional reduction and HDBSCAN clustering”, top) and re-examination (“Re-examination of select cluster(s)”, bottom). **c,** Resulting data from a Method #2 stained HT1080 100K EV sample with non-specific buffer cluster removal. Raw .fcs files from a 100K EV sample and buffer control were analyzed with 50% relative sampling and the stained buffer control cluster was identified (top panels). Re-examination of all remaining clusters, with the exception of the non-specific buffer control cluster, was performed with 80% relative sampling (bottom panels).

**Extended Data Figure 5:**
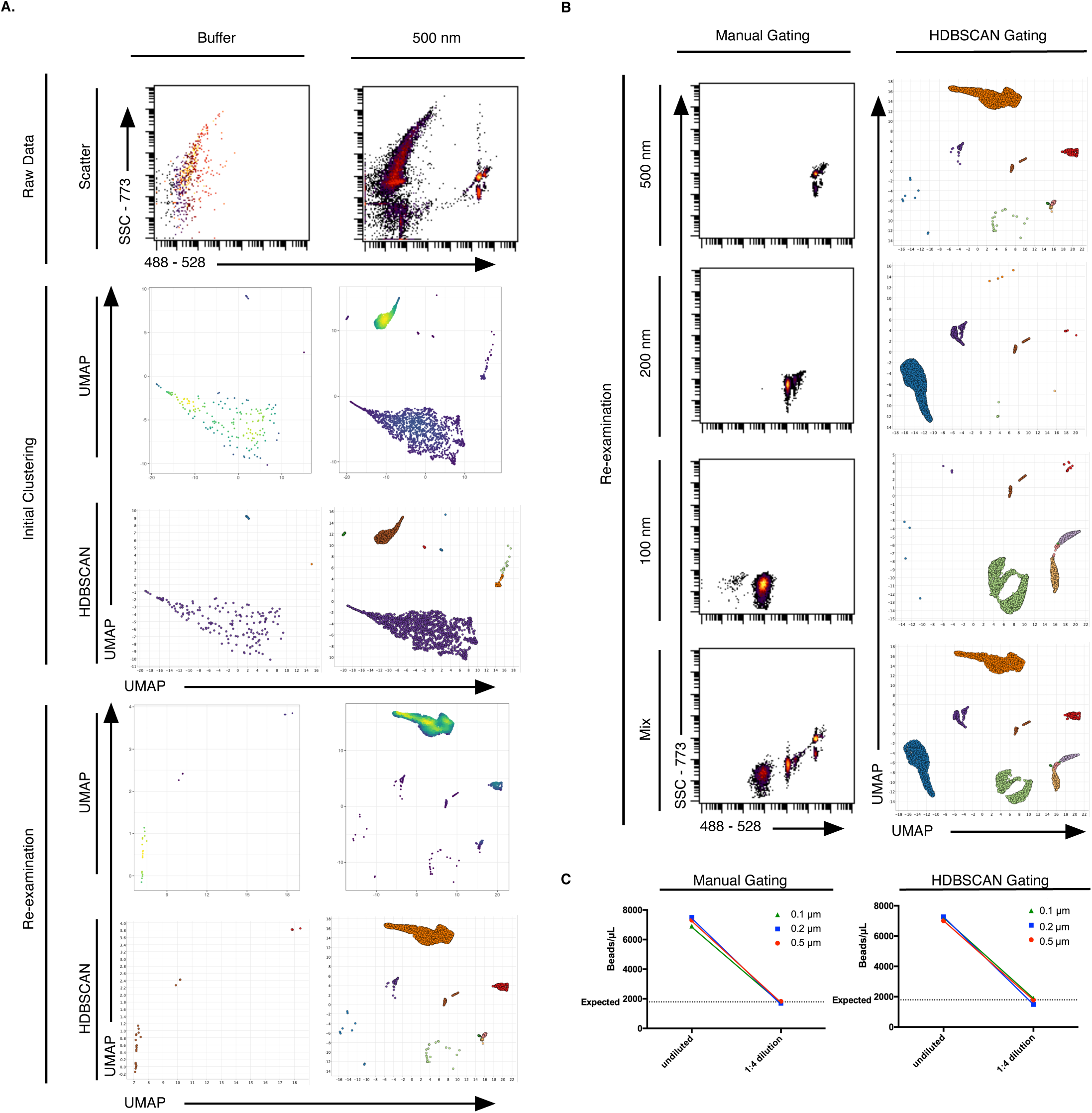
Automated cluster-based gating is concordant with manual gating of synthetic sizing standards. **a,** Strategy for cluster based gating of a non-specific cluster from a blank buffer control. Flow cytometry of FITC labeled sizing beads (500 nm) were acquired with 100% 488-laser power for 2 minutes (top panels). Data were initially analyzed in the EV Fingerprinting pipeline with 80% relative sampling with a non-stained buffer control (“Initial Clustering, middle panels). The buffer-specific cluster was removed and re-examination was performed with 90% relative sampling. **b,** Manual gating (left column) and cluster based HDBSCAN gating (right column) of individual FITC sizing beads (500 nm, 200 nm, and 100 nm) and a mixture (MIX) of all beads. **c,** Quantitative comparison of manual gating (left) and cluster based HDBSCAN gating (right) of FITC sizing beads across two dilutions.

**Extended Data Figure 6:**
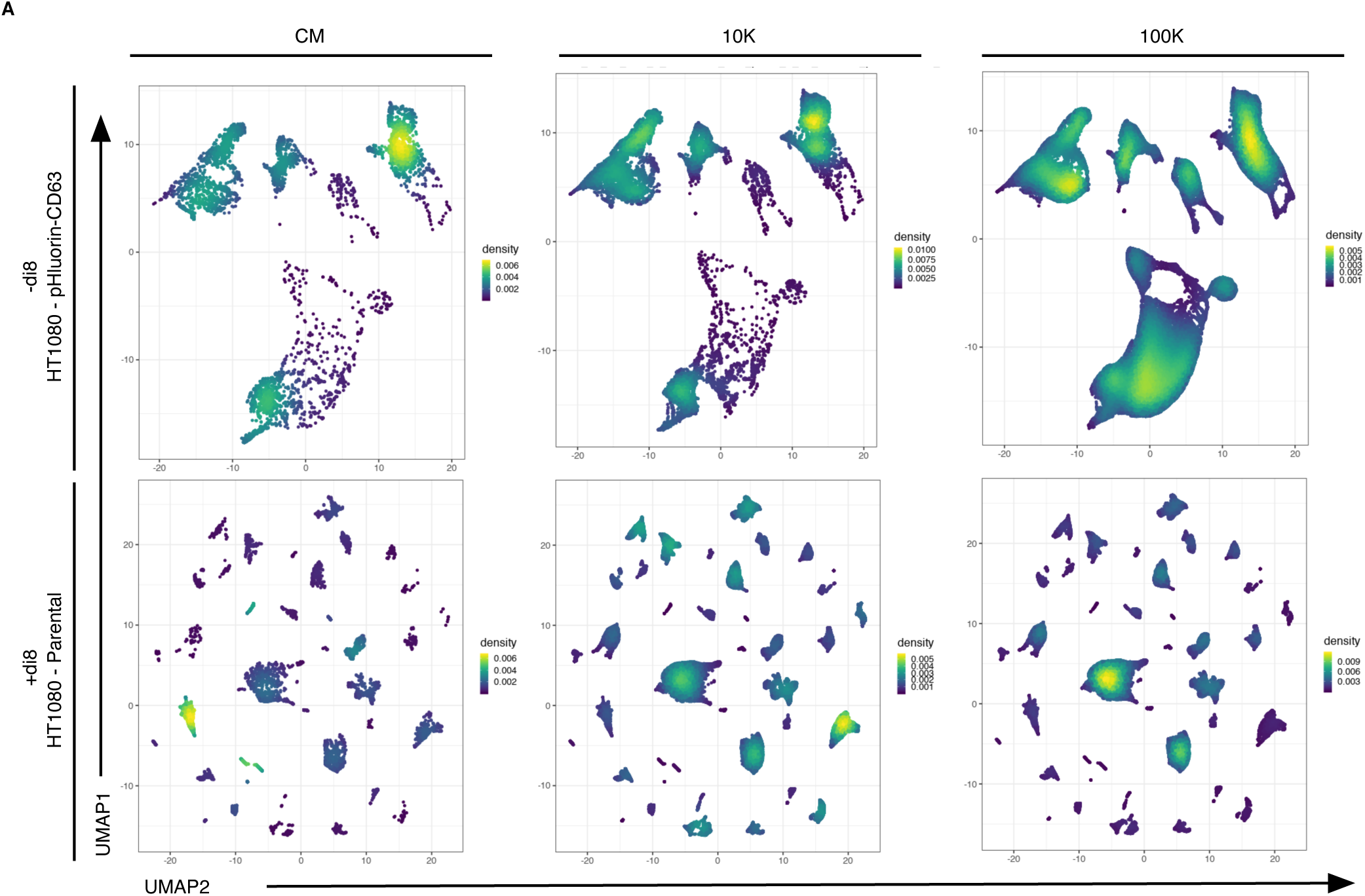
Sample selection criteria fitting for EV Fingerprinting. **a,** Differential enrichment in 10K, 100K, and CM samples visualized using EV Fingerprinting. UMAP embeddings from initial analysis of HT1080-pHluorin-CD63 (top panels, 80% relative sampling) and HT1080-Parental (bottom panels, 50% relative sampling). Dimensional reduction was performed using pHluorin tag and di8 parameters (Supplementary Table 2), respectively.

**Extended Data Figure 7:**
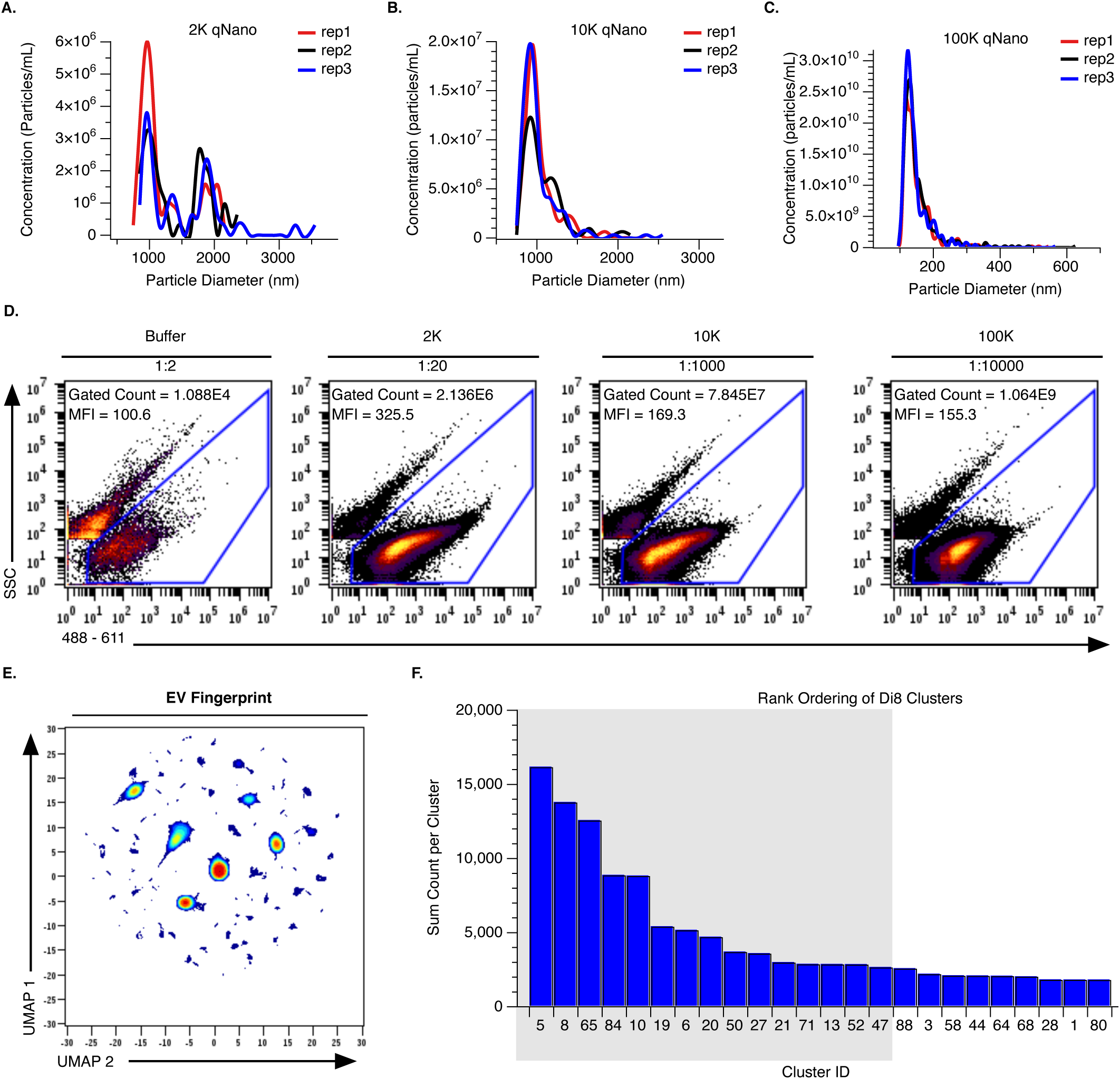
Sizing characterization and staining of DG-UC preps. **a,b,c,** Tunable resistive pulse sensing (TRPS) measurements showing size distribution and particle concentration of density gradient ultracentrifugation (DG-UC) isolated 2.8K (2K, **a**), 10K (**b**), and 100K (**c**) EV from PC3 cells. Each sample measurement was taken as three technical replicates (reps 1-3) after fresh isolation. **d,** Representative flow cytometry scatter plots of blank buffer and DG-UC isolated PC3 EVs (2K, 10K, and 100K) stained with di8 after freeze/thaw at −80℃. 2K and buffer control samples were stained using Method #1 (low particle concentration) while 10K and 100K samples were stained using Method #2 (high particle concentration). Data were collected for 10 min using EV Fingerprinting acquisition settings. Particle positive gating is shown (blue). Gated counts shown are dilution corrected. 488-611 median fluorescence intensities (MFI) by sample are depicted. **e,** Representative EV Fingerprint from initial di8 clustering of buffer controls, 2K, 10K, and 100K DG-UC isolated PC3 EVs. 50% relative sampling for a total of 196,808 events) **f,** Ranked event counts by cluster from initial di8 clustering (**e**) of 2K, 10K, and 100K DG-UC isolated EVs from PC3 cells. Data were sampled relatively (50%, 196,808 events total). Buffer is excluded from total event count. Data shown represent ≥1% of total data. The top 15 clusters (grey box) were selected for re-examination, shown in Figure 3a.

**Extended Data Figure 8:**
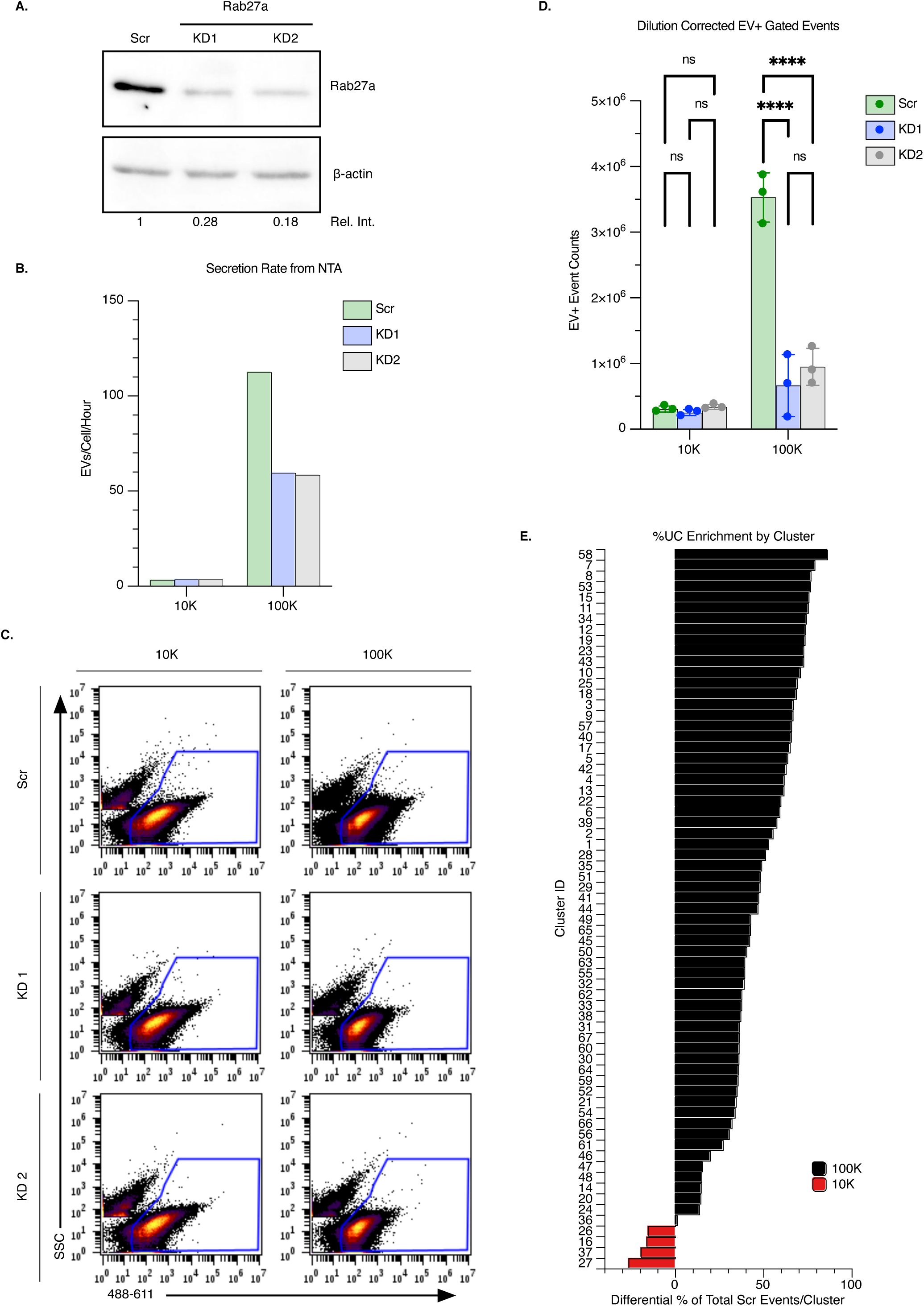
Rab27a KD specifically affects 100K EVs. **a,** Western blotting analysis confirming knock down (KD) of Rab27a protein in HT1080-Scr (Scr) cells compared to HT1080-KD1 (KD1) and HT1080-KD2 (KD2). **b,** EV secretion rate as EVs secreted per cell per hour using nanoparticle tracking analysis (NTA) data from Scr and KD1/KD2 differential ultracentrifugation purified (d-UC) 10K and 100K EV after fresh isolation. **c,** Representative plots depicting flow cytometry gating strategy from Method #2 stained Scr and KD1/KD2 d-UC 10K and 100K EV after storage at 4℃ for less than one week. EV positive (EV+) gating is shown. Data were collected for 10 min using EV Fingerprinting acquisition settings. **d,** Quantitative comparison of EV secretion from 10K and 100K EV from Scr and KD1/KD2 cells by flow cytometry. Technical replicates from three serial dilutions are plotted and EV+ events were corrected for dilution. **** P<0.0001 by two-way ANOVA test. **e,** Enrichment of events per cluster in the 10K or 100K prep quantified as the difference in percent of total from the Scr. Cluster_IDs are represented by decreasing enrichment in the 100K (black) and subsequent increasing enrichment in the 10K (red) from top to bottom.

**Extended Data Figure 9:**
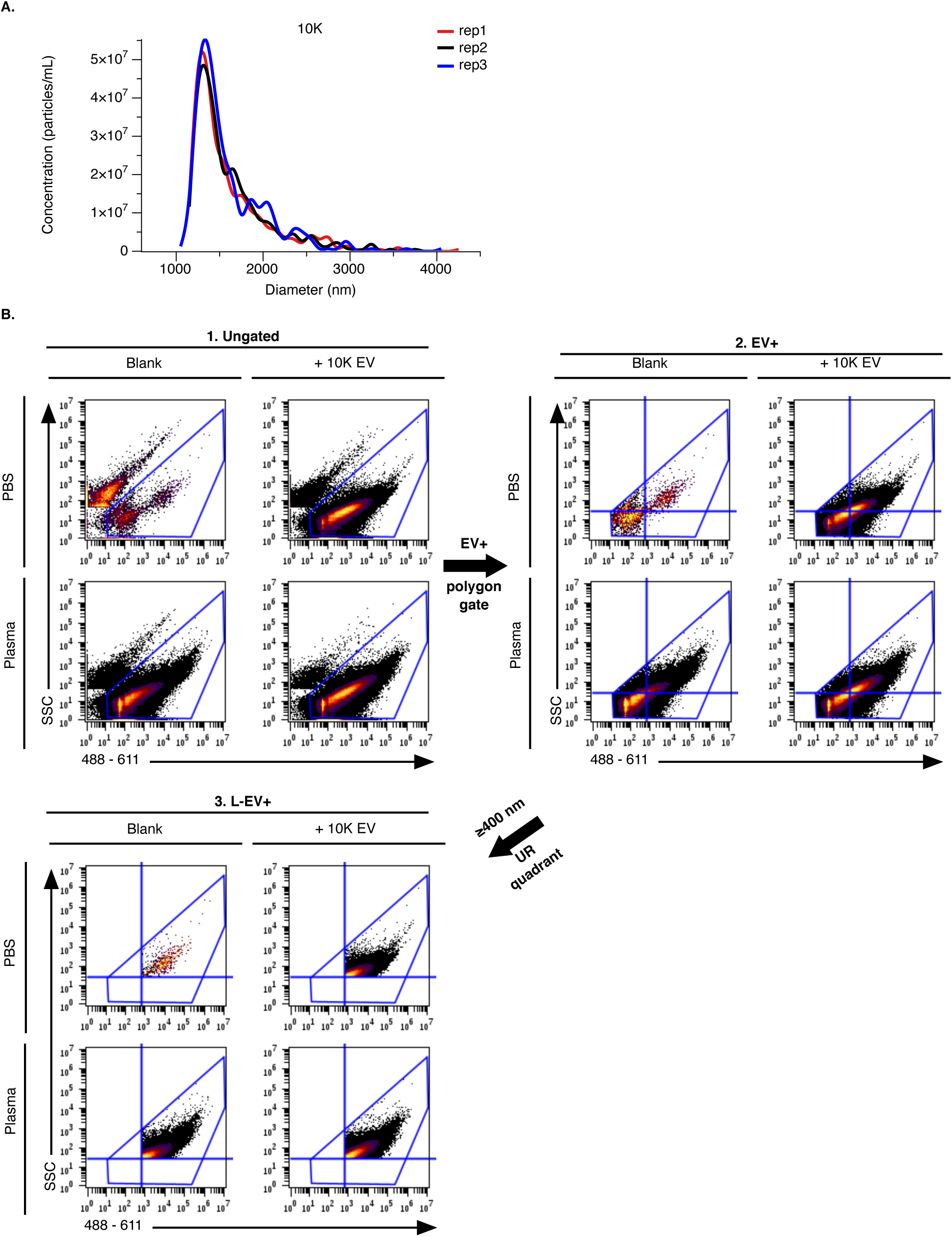
Detection of tumor cell-derived L-EVs in plasma. **a,** Tunable resistive pulse sensing (TRPS) measurements showing size distribution and particle concentration of freeze/thawed (−80℃) differential ultracentrifugation (d-UC) isolated 10K EV from PC3 cells. **b,** Gating strategy for the quantitative detection of L-EV (10K) in plasma by flow cytometry. Ungated data (Step 1) were assessed for EV positive (EV+) events using di8 stained PBS Blank (Step 2, polygon gate). L-EV were determined from di8 staining characteristics (median SSC and 488-611 MFI) of the 400 nm sizing liposomes (from Extended Data Fig. 3 d-f). The upper right quadrant corresponds to EV ≥400 nm (middle panels). Final gated EV+ L-EV+ events (Step 3, quantified in Fig. 6b).

