## Supplementary Table for "EV Fingerprinting: Resolving extracellular vesicle heterogeneity using multi-parametric flow cytometry"

**Supplementary Table 1: List of metrics collected by the CellStream™**

| Index | Feature Name | Feature type | Calculation |
|---|---|---|---|
| 1 | **Camera Timer** | Camera Metric | Digital shutter speed |
| 2 | **Camera Line Number** | Camera Metric | Rate at which exposure and readout can occur |
| 3 | **Time** | Instrument Metric | Time sample was collected, saved in meta data |
| 4 | **FlowSpeed** | Instrument Metric | Fluidics measurement of flow rate, saved in meta data |
| 5 | **Raw Max Pixel [Excitation nm /Emission nm /bandpass nm]** | Optical Metric | Maximum intensity in any pixel without background subtraction |
| 6 | **Raw Min Pixel [Excitation nm /Emission nm /bandpass nm]** | Optical Metric | Minimum intensity in any pixel without background subtraction |
| 7 | **Intensity [Excitation nm /Emission nm /bandpass nm]** | Optical Metric | Sum of all raw pixel intensity with background subtraction |
| 8 | **UCI [Excitation nm /Emission nm /bandpass nm]** | Optical Metric | Uncompensated intensity, same measure as Intensity since we do not perform compensation |
| 9 | **Mean Pixel [Excitation nm /Emission nm /bandpass nm]** | Optical Metric | Mean pixel Intensity after background subtraction |
| 10 | **Bkgd Mean [Excitation nm /Emission nm /bandpass nm]** | Optical Metric | Mean background pixel intensity |
| 11 | **Bkgd StdDev [Excitation nm /Emission nm /bandpass nm]** | Optical Metric | Standard deviation of background intensity |
| 12 | **Area [Excitation nm /Emission nm /bandpass nm]** | Optical Metric | Pixel area of captured signal |
| 13 | **Gradient RMS [Excitation nm /Emission nm /bandpass nm]** | Optical Metric | Mean slope across three pixels, measure of image contrast and focus quality |
| 14 | **Major Axis [Excitation nm /Emission nm /bandpass nm]** | Optical Metric | Longest axis of particle |
| 15 | **Minor Axis [Excitation nm /Emission nm /bandpass nm]** | Optical Metric | Shortest axis of particle |
| 16 | **Aspect Ratio [Excitation nm /Emission nm /bandpass nm]** | Optical Metric | Aspect Ratio of particle |
| 17 | **Saturation Count [Excitation nm /Emission nm /bandpass nm]** | Optical Metric | Number of pixels saturated |
| 18 | **Saturation Percent [Excitation nm /Emission nm /bandpass nm]** | Optical Metric | Percentage of pixels saturated |

**Supplementary Table 2: List of available features and which were used in UMAP and clustering**

| Index | Feature Name | Fluorochrome | Ex (nm) | Calculation | Em (nm) | Used in Initial Analysis? | Used in Re-analysis? |
|---|---|---|---|---|---|---|---|
| 1 | 488 - 528 | Di-8-ANEPPS | 488 | Intensity | 528 | Yes | Yes |
| 2 | RawMaxPixel_488 - 528 | Di-8-ANEPPS | 488 | Raw Max Pixel | 528 | Yes | Yes |
| 3 | Area_488 - 528 | Di-8-ANEPPS | 488 | Area | 528 | Yes | Yes |
| 4 | AspectRatio_488 - 528 | Di-8-ANEPPS | 488 | AspectRatio | 528 | Yes | Yes |
| 5 | 488 - 583 | Di-8-ANEPPS | 488 | Intensity | 583 | Yes | Yes |
| 6 | RawMaxPixel_488 - 583 | Di-8-ANEPPS | 488 | Raw Max Pixel | 583 | Yes | Yes |
| 7 | Area_488 - 583 | Di-8-ANEPPS | 488 | Area | 583 | Yes | Yes |
| 8 | AspectRatio_488 - 583 | Di-8-ANEPPS | 488 | AspectRatio | 583 | Yes | Yes |
| 9 | 488 - 611 | Di-8-ANEPPS | 488 | Intensity | 611 | Yes | Yes |
| 10 | RawMaxPixel_488 - 611 | Di-8-ANEPPS | 488 | Raw Max Pixel | 611 | Yes | Yes |
| 11 | Area_488 - 611 | Di-8-ANEPPS | 488 | Area | 611 | Yes | Yes |
| 12 | AspectRatio_488 - 611 | Di-8-ANEPPS | 488 | AspectRatio | 611 | Yes | Yes |
| 13 | 488 - 702 | Di-8-ANEPPS | 488 | Intensity | 702 | Yes | Yes |
| 14 | RawMaxPixel_488 - 702 | Di-8-ANEPPS | 488 | Raw Max Pixel | 702 | Yes | Yes |
| 15 | Area_488 - 702 | Di-8-ANEPPS | 488 | Area | 702 | Yes | Yes |
| 16 | AspectRatio_488 - 702 | Di-8-ANEPPS | 488 | AspectRatio | 702 | Yes | Yes |
| 17 | 488 - 773 | Di-8-ANEPPS | 488 | Intensity | 773 | Yes | Yes |
| 18 | RawMaxPixel_488 - 773 | Di-8-ANEPPS | 488 | Raw Max Pixel | 773 | Yes | Yes |
| 19 | Area_488 - 773 | Di-8-ANEPPS | 488 | Area | 773 | Yes | Yes |
| 20 | AspectRatio_488 - 773 | Di-8-ANEPPS | 488 | AspectRatio | 773 | Yes | Yes |
| 21 | 642-702 | anti-CD63-APC | 642 | Intensity | 702 | No | Yes |
| 22 | RawMaxPixel_642-702 | anti-CD63-APC | 642 | Raw Max Pixel | 702 | No | Yes |
| 23 | Area_642-702 | anti-CD63-APC | 642 | Area | 702 | No | Yes |
| 24 | AspectRatio_642-702 | anti-CD63-APC | 642 | AspectRatio | 702 | No | Yes |
| 25 | 488 - 528 | CD63-pHluorin | 488 | Intensity | 528 | Yes | Yes |
| 26 | RawMaxPixel_488 - 528 | CD63-pHluorin | 488 | Raw Max Pixel | 528 | Yes | Yes |
| 27 | Area_488 - 528 | CD63-pHluorin | 488 | Area | 528 | Yes | Yes |
| 28 | AspectRatio_488 - 528 | CD63-pHluorin | 488 | AspectRatio | 528 | Yes | Yes |

**Supplementary Table 3: Summary of staining method conditions**

| Method: | Sample Concentration: | Di8 Staining Concentration (μM): | 200x Post-Stain dilution? | Final Di8 Concentration: | Staining Conditions: | MC Buffer Stock: | Laser Power: |
|---|---|---|---|---|---|---|---|
| 1 | Low (Ex: CM, dilute EV preps) | 0.25 μM | no | 0.25 μM | 1 hr RT | 6.5% | 25% |
| 2 | High (Ex: 10K and 100K EVs, plasma) | 2 μM | yes | 0.01 μM | 1 hr (-ab)<br>3 hr (+ab)<br>Both @ 37 C | 6.5% | 25% |

**Supplementary Table 4: Sizing liposome standards composition and size**

| Sample | Composition | Lipid Ratios | Z-Maximum (nm) |
|---|---|---|---|
| 400 (SUVs) | DSPC/Chol/DMG-$PEG_{2000}$ | (52:45:3) | 458.67 |
| 200 (SUVs) | DSPC/Chol/DMG-$PEG_{2000}$ | (52:45:3) | 190.14 |
| 90 (SUVs) | DSPC/Chol/DMG-$PEG_{2000}$ | (52:45:3) | 91.28 |

**Supplementary Table 5: Cholesterol liposome standards composition and size**

| Sample | Composition | Lipid Ratios | Z-Maximum (nm) |
|---|---|---|---|
| 0% Cholesterol | DSPC/DMG-$PEG_{2000}$ | (95:5) | 105.71 |
| 10% Cholesterol | DSPC/Chol/DMG-$PEG_{2000}$ | (87:10:3) | 122.42 |
| 25% Cholesterol | DSPC/Chol/DMG-$PEG_{2000}$ | (72:25:3) | 105.71 |
| 45% Cholesterol | DSPC/Chol/DMG-$PEG_{2000}$ | (52:45:3) | 122.42 |

**Supplementary Table 6: EV Fingerprinting pipeline clustering parameters**

| Clustering Parameter (“set_parameters” node): | HDBSCAN min_cluster size | UMAP n_neighbors | UMAP n_epochs | HDBSCAN min_samples | UMAP min_dist |
|---|---|---|---|---|---|
| Set Value: | 100 | 15 | 1,000 | 200 | 0.1 |
